# Heterogeneous dynamics of mobile genetic elements encode and spread antibiotic persistence in bacteria

**DOI:** 10.64898/2026.02.27.708469

**Authors:** Iris Dadole, Romain Gory, Kevin T. Huguet, Natacha Lenuzza, Friederike Steurer, Benoît Morin, Christophe Ginevra, Laetitia Attaiech, Xavier Charpentier, Sophie Jarraud, Didier Blaha, Alexander Schmidt, Jean-Philippe Rasigade, Nicolas Personnic

## Abstract

Bacterial persisters are nonreplicating cells that transiently evade antibiotic killing within an otherwise susceptible, replicating population. While persister formation has traditionally been attributed to phenotypic heterogeneity, reversible functional variations among genetically identical cells, most studies lack direct genomic evidence to exclude the contribution of cryptic and reversible genetic variations. Here, we demonstrate that the dynamics of the *Legionella pneumophila* integrative and conjugative element (ICEs) pP36 encode and disseminate antibiotic persistence-like traits. During biofilm formation, pP36 undergoes heterogeneous chromosomal excision generating a subpopulation of nonreplicating persisters-like bacteria with a distinct molecular profile. pP36-mediated *L. pneumophila* growth rate heterogeneity significantly enhances bacterial survival under fluoroquinolone treatment. Our findings reveal that pP36 dynamics not only orchestrate physiological diversity but also serve as a vehicle for horizontal transfer of antibiotic persistence mechanisms among clinical *Legionella* isolates. While ICEs are recognized as key drivers of antibiotic resistance dissemination, this study establishes that ICEs-mediated phenotypic variations is a critical determinant of persister-like cell formation, reshaping our understanding of bacterial adaptability to antibiotic stress.

## Main

Bacterial communities exhibit a remarkable ability to adapt to complex and changing environments through functional specialization among their members, within the micro-meter range ^1–4^. A primary example is the bacterial persisters. A persister is a nonreplicating individual bacterium, within an otherwise proliferative population, that transiently evades the bactericidal activity of antimicrobials ^5^. Persisters are a major driver of antibiotic treatment failure and infection recurrence, posing a critical public health challenge ^6^. The formation of persisters has been experimentally documented for many major bacterial pathogens including *Staphylococcus aureus* ^7–9^, *Mycobacterium tuberculosis* ^10,11^, *Escherichia coli* ^12,13^, *Salmonella enterica* ^14,15^, *Pseudomonas spp*.^16^, *Listeria monocytogenes* ^17^, *Legionella pneumophila* ^18^, *Burkholderia pseudomallei* ^19^, *Yersinia pseudo-tuberculosis* ^20^, *Streptococcus pneumoniae* ^21^, *Brucella abortus* ^22^ or *Klebsiella pneumoniae* ^23^.

Persister formation has traditionally been attributed to phenotypic heterogeneity, *i.e.*, reversible functional variations among genetically identical cells ^24–26^. Yet, most studies lack direct genomic evidence to exclude the contribution of structural genetic variations such as those mediated by mobile genetic elements (MGEs). MGEs represent a broad family of DNA sequences that can move or be transferred between different locations within a genome (excision, integration, transposition, replication) or between genomes (conjugation, transduction, vesiduction or transformation) ^27–31^. Integrative and conjugative elements (ICEs) represent the most prevalent type of conjugative element ^32,33^. ICEs can: (1) integrate into and excise from host chromosomes at specific sites, (2) form circular dsDNA intermediates capable of rolling-circle replication and horizontal dissemination via conjugation, and (3) carry adaptive gene cargo, including antibiotic and heavy metal resistance genes, metabolic pathways for alternative carbon sources, and virulence or symbiosis factors ^34–37^. The extent to which phenotypic variations and persister-like cells formation within a seemingly identical population arise from ICE-driven reversible structural variations of the genome remains unknown.

*L. pneumophila*, the bacterium responsible for Legionnaires’ disease ^38^, provides an ideal system to study how ICEs influence antibiotic persistence. First, its genome is highly shaped by mobile genetic elements, including genomic islands (GIs) and ICEs, offering a natural framework to investigate their role in bacterial adaptability ^39–47^. Second, our previous work has shown that *L. pneumophila* exhibits marked growth rate heterogeneity during biofilm and intracellular growth, resulting in coexisting subpopulations of actively dividing individuals and nonreplicating antibiotic persisters ^18,48,49^.

While persister formation has long been attributed to stochastic noise in cellular circuits, we uncover a hidden genomic dimension to this phenomenon. Here, we reveal that the integrative and conjugative element pP36 acts as a molecular switch, dynamically reshaping *L. pneumophila* populations by generating growth rate heterogeneity and promoting the emergence of nonreplicating antibiotic persister-like cells. Crucially, pP36 is not just a passive player as it may hijack host regulatory networks via a *csrA*-like cargo gene, and spreads these traits horizontally among clinical isolates. Our findings expose mobile genetic elements as architects of phenotypic diversity, offering a radical shift in how we understand, and potentially combat, antibiotic tolerance. By targeting the dynamic interplay between ICEs and host physiology, we may finally disrupt the survival strategies that allow pathogens to evade drugs.

## Results

### Stochastic ICE dynamics favors bacterial growth rate heterogeneity and antibiotic persistence

To question the interaction between ICE dynamics and antibiotic persistence, we developed a computational simulation framework to model how dynamic excision and integration of ICEs influence population-level diversity and the formation of an antibiotic-tolerant subpopulation in *L. pneumophila*. We first sketched a qualitative, mechanistic account of how ICE mobility couples to fluoroquinolone action to generate biphasic time-kill curves, a hallmark of bacterial persistence. The assumption that fluoroquinolone lethality has a growth- or metabolism-dependent component is consistent with experimental literature showing reduced killing in nongrowing *L. pneumophila* populations ^18,49^ and other species ^50–54^. We used the *msevol* stochastic framework (https://github.com/rasigadelab/msevol), in an explanatory rather than calibrative mode, to explore whether a small set of biological rules was sufficient to reproduce biphasic time-kill curves, a hallmark of bacterial antibiotic persistence (**Fig. 1; Fig. SI 1 to 7; SI Table 1, 2**). We began with a homogeneous, growing population under a fluoroquinolone regimen and observed a single, steep survival decline without a late tail (**Fig. 1a**). This baseline confirms that, in the absence of growth-rate heterogeneity, the model does not generate biphasic time-kill curves. When we introduced intrinsic growth variation independent of ICE dynamics, survival curves showed a modest curvature (**Fig. 1b; Fig. SI 8, 9**). By introducing stochastic ICE excision, which abruptly slows growth, we observed a split into two populations under antibiotic treatment: a rapidly killed, dividing majority and a slowly killed, ICE-excised subpopulation, producing a distinct biphasic survival curve (**Fig. 1c; Fig. SI 10, 11**). Blocking the ICE excision *in silico* abolished the slow-killing phase underlying its importance for antibiotic persistence (**Fig. SI 10, 11**). Furthermore, enabling ICE autonomous replication in a subset of excised ICEs prolonged the survival tail (minimum duration of killing, MDK) without altering minimal inhibitory concentration (MIC), a signature of antibiotic persistence (**Fig. 1d; Extended Data Fig. 1; Fig. SI 10, 11**). Guided by our simulation model, we propose that stochastic ICE mobility, cycling between integrated, excised, and replicating states, imposes growth penalties on a subset of *L. pneumophila* individuals forming a persister-like subpopulation that enhance population survival under fluoroquinolones exposure. We translated this model into two experimental questions and addressed them with matched assays: Does ICE dynamics correlate with *L. pneumophila* growth rates? Does ICE excision/replication control non-grower formation and fluoroquinolone persistence?

**Figure 1:**
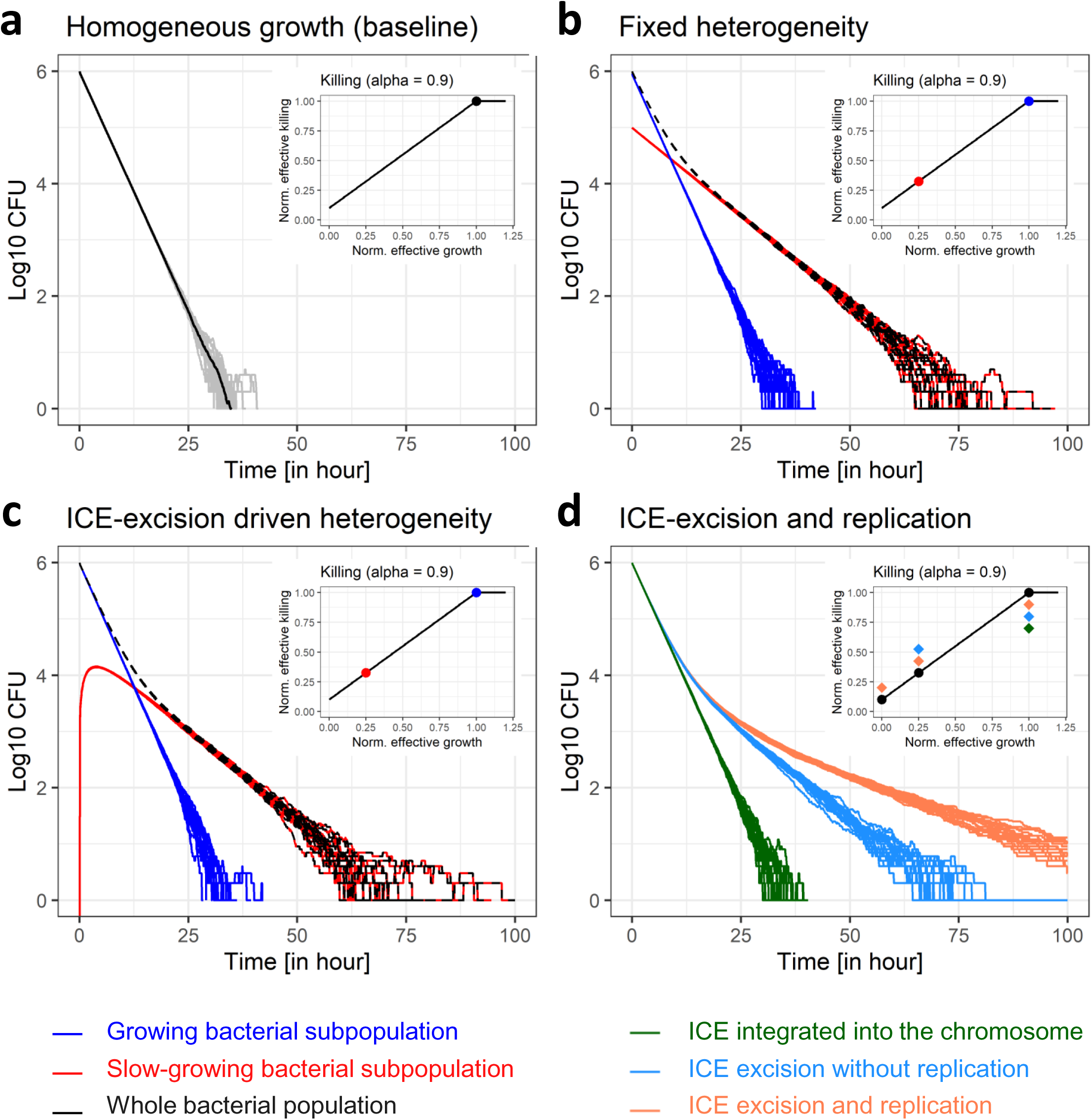
Stochastic ICE dynamics favors bacterial growth rate heterogeneity and antibiotic persistence. Simulated time-kill assays of bacterial cell populations under constant antibiotic pressure with different sources of growth heterogeneity. Simulation parameters common to all panels are listed in supplementary information Table 2. Panels **a-c** display the size of the total bacterial population over time (black curves). Depending on the model, this total population may consist of distinct subpopulations, shown as colored curves. Panel **d** displays only the total population dynamics for several alternative models corresponding to the most comprehensive ICE framework (*i.e*., allowing ICE excision and intracellular replication), with the aim of comparing how individual ICE behaviors affect the survival dynamics of the whole cell population. Although cell diversification occurs in some of these models, the resulting subpopulations are not explicitly shown (see examples in Extended Data Fig. 1). In this panel, curve colors denote the ICE behaviors allowed in each model rather than individual cell subpopulations. (**a**) Baseline simulated time-kill curves for a homogeneous cell population growing at its intrinsic growth rate. By construction, no cell diversification occurs. Grey curves show *N* = 20 simulation replicates, while the black curve represents the corresponding average dynamics. A monophasic exponential decline, appearing as a straight line on the log_10_ scale, is observed with all cells experiencing maximal killing. (**b**) Simulated time-kill curve for an initially heterogeneous cell population composed of two subpopulations growing at different rates (*N* = 20 replicates). The black curve represents the total population, defined as the sum of a normally growing subpopulation (blue) and a slow-growing subpopulation (red), the latter initially comprising 10% of the total population. Under strong growth–killing coupling (α = 0.9), slow-growing cells survive longer, generating a detectable inflection in the total population trajectory and resulting in a biphasic time-kill profile. **(c)** Simulated time-kill curve for an initially homogeneous population diversifying through integrative and conjugative element (ICE) excision (*N* = 20 replicates). The total population (black) consists of two subpopulations: cells carrying a silent, integrated ICE and growing at their intrinsic rate (blue), and cells in which ICE excision has occurred, leading to an abrupt and irreversible growth slowdown (red). To test whether ICE excision alone can generate sufficient growth heterogeneity to produce biphasic killing (as in panel B), we considered a scenario in which a fraction of cells randomly excises the ICE, triggering an immediate and irreversible reduction of growth. The ICE-associated fitness cost is expressed only upon excision, and excised cells maintain a constant reduced growth rate throughout the simulation, with no further diversification. Simulations were performed for high growth-killing coupling parameter (α = 0.9), with an excision rate of 0.01 per hour (compatible with switching values estimated by other modeling approaches ^94^). A biphasic profile, with a longer tail persistence is noticeable. (**d**) Comparison of simulated time-kill curves for an initially homogeneous population diversifying through different integrative and conjugative element (ICE) behaviors. Curves represent the total cell population (*N* = 20 simulations for each model). The color of each curve denotes the ICE behavior allowed in the corresponding model (not individual cell subpopulations). Dark green: ICE cannot excise, preventing any diversification from the initial homogeneous genotype (similar to panel a). Light blue: ICE can excise, generating growth heterogeneity through an abrupt reduction of growth in excised cells (similar to panel c). Coral: ICE can excise and modulate its copy number, producing further diversification with variable growth rates depending on ICE copy number (as illustrated in Extended Data Fig. 1). This panel highlights how different ICE behaviors can generate distinct population-level killing dynamics: no excision results in a monophasic decline, excision alone generates limited growth heterogeneity and a delayed biphasic response, while excision combined with copy number modulation produces pronounced subpopulation diversity and a clearer biphasic profile. Note that each panel (a-d) includes an inset showing the apparent antibiotic-mediated killing as a function of effective bacterial growth. Points represent the growth and killing characteristics of cells either initially present or generated through ICE dynamics. In panels a-c, these points are colored according to their effective growth rate, matching the colors used in the main plots. In panel d, points are colored according to the ICE model in which the corresponding subpopulation behavior is observed (*i.e*., a point matching the color of the main plot is added whenever that ICE model produces a subpopulation with the corresponding growth/killing characteristics).

### pP36 ICE dynamics is heterogeneous and bound to the growth status of sessile *L. pneumophila*

We experimentally tested our simulation-based hypothesis during biofilm growth, conditions known to induce growth rate heterogeneity in *L. pneumophila* and antibiotic persistence ^49^. We monitored the bacterial growth dynamics by using the Timer single-cell growth rate reporter ^15^ where the green-to-red Timer fluorescence ratio (Log_10_[500 nm/600 nm]) reflects defined *L. pneumophila* single-cell growth rates, as previously described in ^18, 49^. Green-dominant cells are actively dividing, while red-dominant cells are growth-arrested. During biofilm formation, Timer producing *L. pneumophila*, showed various fluorescent ratio (R) reflecting different bacterial division rates, including a fraction of growth-arrested individuals (R < 0; growth rate [μ] = 0.0 h^−1^) (**Fig. 2a-b; Extended Data Fig. 2a**). To explore a potential role of ICEs in the observed bacterial growth rates variations, we first screened for activity the five ICEs or GIs of *L. pneumophila* Paris, LppGI-A (GI, 12.2 kb), pP36 (ICE, 37.3 kb), LppGI-1 (ICE, 44.6 kb), LppGI-B (GI, 11.7 kb), and LppGI-2 (ICE, 123.9 kb) (**Extended Data Fig. 2b**), by endpoint PCR targeting circular intermediates (**Fig. 2c, Extended Data Fig. 2c**). Only LppGI-A and pP36 showed detectable activity, with pP36 identified across most conditions. Next, we quantified the subpopulation-level dynamics of LppGI-A and pP36. Using quantitative PCR (qPCR), we measured the frequency of empty integration sites in approximately 10⁶ growers and non-growers, which had been isolated by FACS based on their Timer fluorescence profiles (**Fig. 2b, 2d**). Strikingly, pP36 exhibited ∼100-fold higher excision activity in non-proliferating bacteria compared to proliferating cells (**Fig. 2d**). This pattern was also observed in slow-growing bacteria (0 < R < 0.2; μ = 0.04 h⁻¹), implicating that both reduced growth rates and growth-arrest correlate with elevated ICE activity (**Fig. 2b; Extended Data Fig. 2d**). In contrast, LppGI-A showed no growth-dependent excision pattern. Together, these findings show that the behaviour of the ICE pP36 is heterogeneous in biofilms, and that its dynamics are correlated with the growth rate of individual cells.

**Figure 2:**
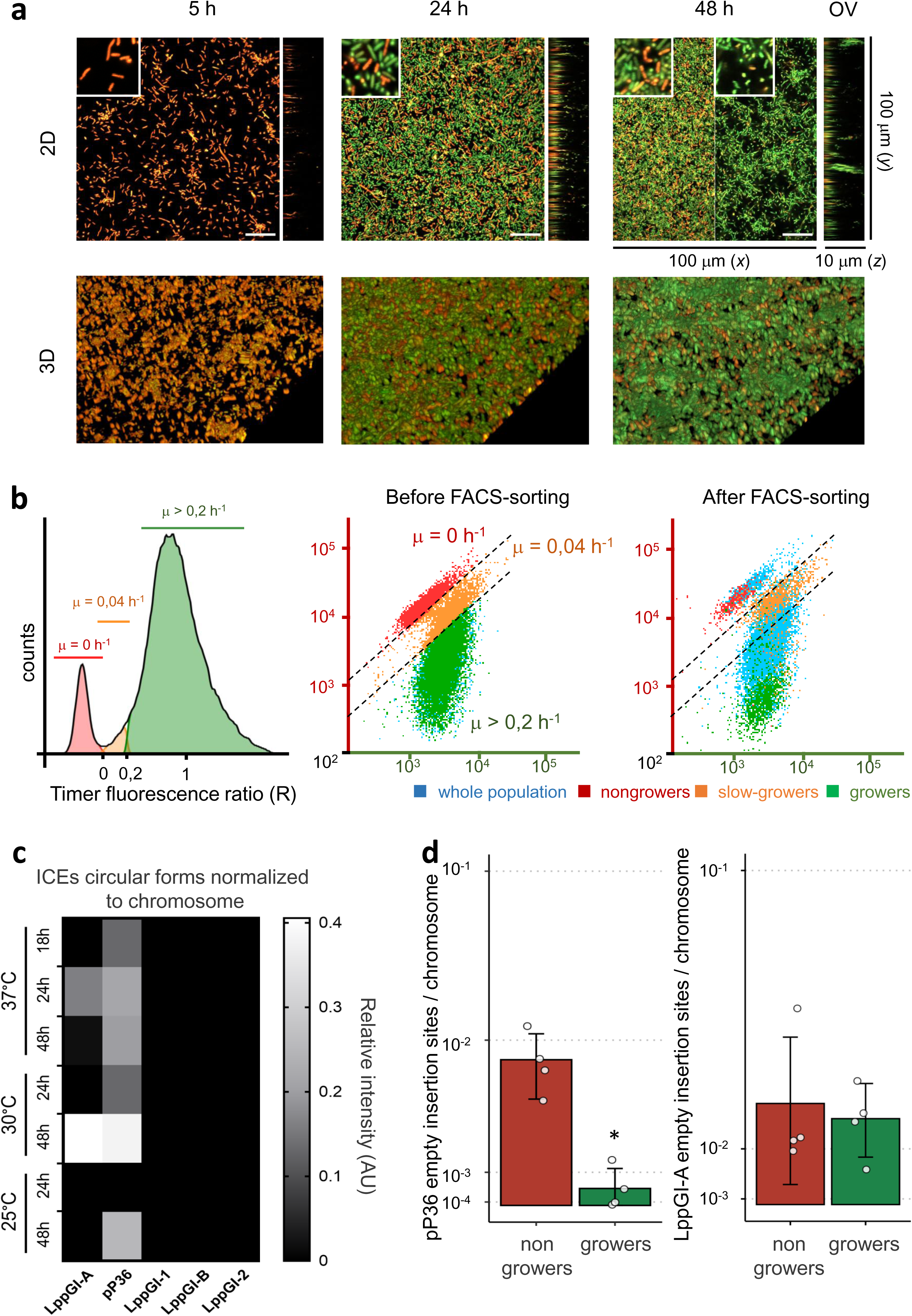
pP36 ICE dynamics is heterogeneous and bound to *L. pneumophila* growth status. *L. pneumophila* producing Timer (pNP107) were grown to stationary phase, diluted in AYE broth, and allowed to form biofilms at 30 °C. (**a**) Confocal microscopy analysis. Upper panel: 2D micrographs show overlays of the Timer fluorescence (500 nm and 600 nm); growing and nongrowing bacteria appear green or red/orange, respectively. OV: orthogonal view. Scale bars 10 μm. Lower panel: 3D reconstructions (x = 100 μm, y = 100 μm, z = 10 μm). (**b**) Biofilms formed after 48 h were homogenized and analysed by flow cytometry. The Timer fluorescence (500nm and 600nm) was recorded for individual bacterium. Left panel: Frequency graph showing the distribution of the Timer fluorescence ratio (Log_10_[500 nm/600 nm]). Corresponding bacterial division rates are indicated according to ^49^. Central panel: Dot plot showing Timer spectral properties for individual bacterium and the corresponding division rates (μ). Left panel: Dot plot showing the FACS-sorting of intracellular *L. pneumophila* subpopulations. Bacteria were FACS-sorted according to their Timer spectral properties and corresponding division rate. Sorted subpopulations were re-analysed by flow cytometry and compared with the initial population to evaluate the separation efficiency. (**c**) PCR-based heatmap screening of five GIs and ICEs in circular form in biofilms grown at different temperatures (25°C, 30°C, and 37°C) and growth times (18 h, 24 h, and 48 h). Relative PCR band intensity is shown normalized to chromosomal DNA. (**d**) qPCR analysis of bacterial subpopulations from 48 h biofilms grown at 30°C. *L. pneumophila* were sorted based on Timer fluorescence properties, genomic DNA extracted and pP36 and LppGI-A empty insertion site quantified by qPCR. Data represent the mean ± SD of four biological replicates (two-tailed Student’s t test; \*\*\**P* < 0.001, ** *P* < 0.01, \**P* < 0.05).

### pP36 ICE dynamics orchestrates sessile *L. pneumophila* growth rate heterogeneity

To directly assess the role of ICEs in generating heterogeneity in bacterial growth rates, we first disrupted pP36 mobility by deleting the integrase/excisionase gene *lpp0194*. In the resulting *L. pneumophila* Δ*int* mutant, pP36 excision was abolished, as confirmed by qPCR analysis of biofilms (**Extended Data Fig. 3a-b**). We then evaluated the distribution of *L. pneumophila* growers and nongrowers, as defined by the Timer color ratio, by flow cytometry analysis of homogenized biofilms. It revealed a marked 10-fold decrease in nongrowing subpopulation size in the Δ*int* mutant as compared to the parental strain (**Fig. 3a-b**). Concomitantly, disrupting pP36 excision increased the growth rate of dividing cells (**Fig. 3a**; median growth rate: 0.2 h⁻¹ in parental vs. 0.3 h⁻¹ in Δ*int*, as indicated by Timer fluorescence ratios of 0.3 and 0.5, respectively, according to ^18^). To map the spatial distribution of growing and nongrowing cells within the *L. pneumophila* biofilm architecture, we performed confocal microscopy on biofilms and processed the images using our in-house pipeline for single-cell growth rate tracking (**Fig. 3c-d; Extended Data Fig. 4a**). In the parental strain, nongrowers were dispersed throughout the biofilm but enriched in the basal layer, while growers dominated the upper layers. Disrupting pP36 dynamics (Δ*int* mutant) reduced non-growers across all biofilm layers. To determine whether nongrowers represent cells that fail to initiate growth during biofilm formation, we immobilized in agarose individual Timer-producing parental and Δ*int* strains in imaging chambers ^55^ and monitored microcolony formation via confocal microscopy ^49,55^ (**Fig. 4a-b; Extended Data Fig. 4b**). While about 30% of sessile wild-type cells grew and form green microcolonies, 75% of Δ*int* individuals did so. The mathematical model (**Fig. 1c-d; Fig. SI 10, 11**) suggested that ICEs autonomous replication would contribute to the formation of growth-arrested bacteria. We thus inactivated pP36 origin of replication and abolished circular pP36 forms in the resulting Δ*oriV* mutant (**Extended Data Fig. 3b**). This disruption resulted in a decrease in the nongrowing subpopulation size in biofilms (**Fig. 3a-d; Extended Data Fig. 4a**), accompanied by increased growth resumption (**Fig. 4a-b**). In summary, pP36 ICE dynamics contribute to *L. pneumophila* phenotypic noise, namely the coexistence of growing and growth-arrested *L. pneumophila* in biofilms, likely through regulation of growth initiation on surfaces.

**Figure 3:**
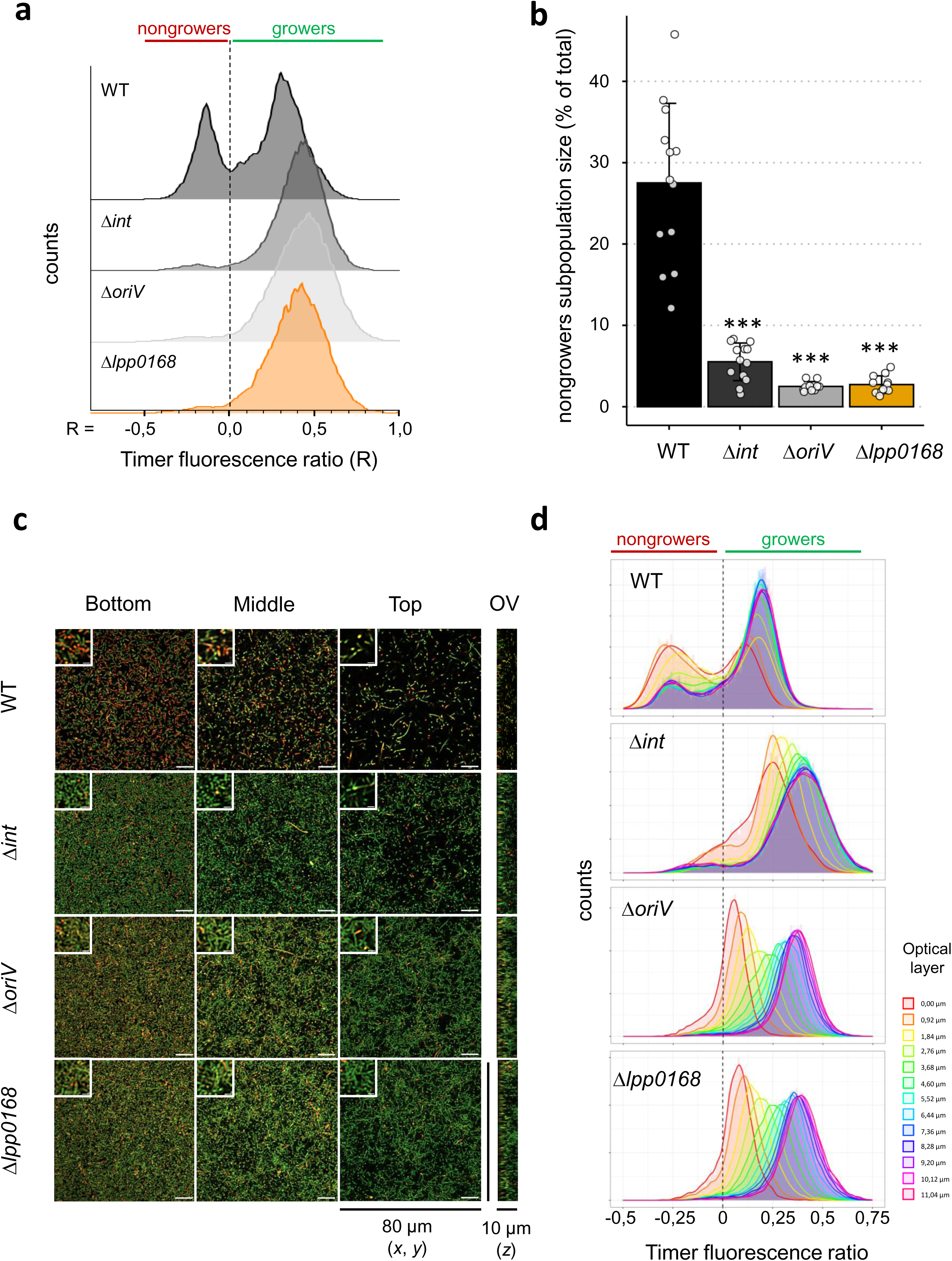
pP36 ICE dynamics orchestrates sessile *L. pneumophila* growth rate heterogeneity. *L. pneumophila* wild-type (WT) and the isogenic mutants Δ*int*, Δ*oriV* and Δ*lpp0168*, producing Timer, were grown to stationary phase, diluted in AYE broth, and allowed to form biofilms at 30 °C for 24 h. (**a-b**) Biofilms were homogenized and analysed by flow cytometry. The Timer fluorescence (500nm and 600nm) was recorded for individual bacterium. (a) Frequency graph showing the distribution of the Timer fluorescence ratio (Log_10_[500 nm/600 nm]). (b) Quantification of the subpopulation size in nongrowers. Data represent the mean ± SD of six biological replicates (two-tailed Student’s t test; \*\*\**P* < 0.001, ** *P* < 0.01, \**P* < 0.05). (**c-d**) Confocal microscopy analysis. (c) Micrographs (*x* = 80 μm, *y* = 80 μm) show overlays of the Timer fluorescence (500 nm and 600 nm); growing and nongrowing bacteria appear green or red/orange, respectively. OV: orthogonal view (*z* = 10 μm). Scale bars 10 μm. (d) Distribution of the Timer fluorescence ratio calculated for each individual bacterium at different optical layer in the biofilm.

**Figure 4:**
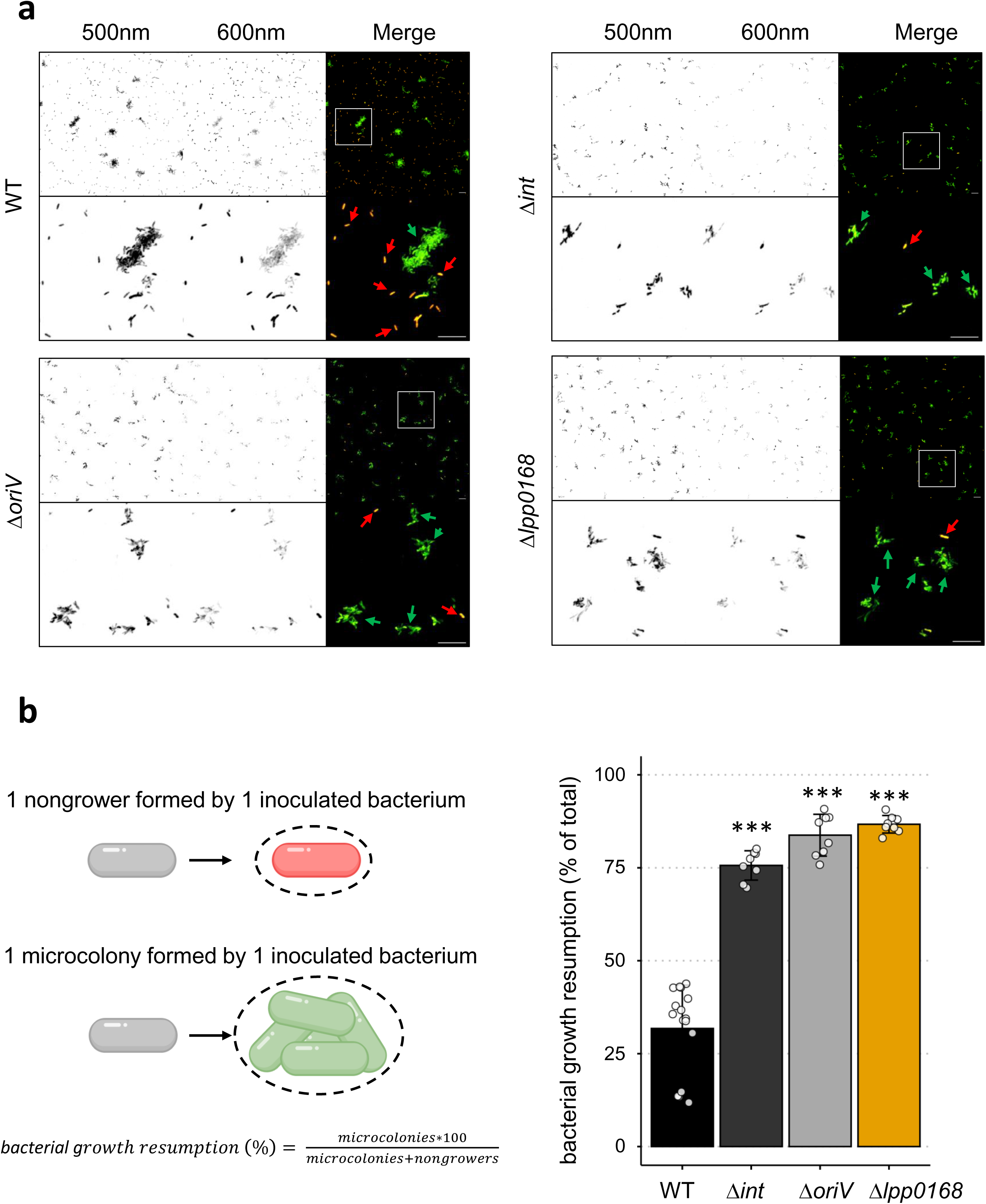
pP36 control growth resumption of sessile *L. pneumophila*. Stationary phase Timer-producing *L. pneumophila* wild-type (WT) and the isogenic mutants Δ*int*, Δ*oriV* and Δ*lpp0168* were immobilized in AYE/0.5% agarose, and let form microcolony for 48 h. (**a**) Fluorescence micrographs of microcolonies (scale bars, 20 μm). Magnification are showed. Red arrow: Nongrowing cells. Green arrow: Proliferating cells forming microcolonies. (**c**) Quantification of bacterial growth resumption. Data represent the mean ± SD of a minimum of four biological replicates (two-tailed Student’s t test; \*\*\**P* < 0.001, ** *P* < 0.01, \**P* < 0.05).

### pP36 ICE dynamics shapes the molecular atlas of *L. pneumophila* individuals with reduced growth

Given that pP36 dynamics correlate with reduced growth rate (**Fig. 3; Extended Data Fig. 2d**), we hypothesized that this ICE might reprogram the molecular identity of non/slow-growers to maintain their state. Comparative proteomics of FACS-sorted non/slow-growers from wild-type and Δ*int* biofilms identified 2,149 proteins, of which only 61 were specifically enriched in wild-type non/slow-growers, compared to only four in Δ*int*. This suggested that pP36 actively reshaped their molecular profile (**Fig. 5a,c; Extended Data Table 1**). This reprogramming appears specific to non/slow-growers, as pP36 mobility had minimal effects on growing cells (**Fig. 5b; Extended Data Table 2**), aligning with its 100-fold higher excision activity in these subpopulations (**Extended Data Fig. 2d**). The 61 proteins spanned several key functional categories (**Fig. 5c; Extended Data Fig. 5; Extended Data Table 1**). It included: (i) respiratory and redox capacity (NADH dehydrogenase, succinate dehydrogenase (SdhB/D), malate dehydrogenase (MaeA), and short-chain dehydrogenases); (ii) metabolism, carbon storage and lipids utilization (NfeD, polyhydroxyalkanoate synthetase, lipases, and esterases); (iii) stress response (acid-resistance protein HdeD, Na⁺/H⁺ antiporters, and the multidrug efflux pump HylD, detoxifying enzyme AhpD); (iv) virulence and motility (flagellin, MotA, PilY1, PilZ, and components of the type 2 and 4 secretion systems), (v) cell envelop factors (Lipopolysaccharide Transport Protein C, FtsW, Small LPS-binding Proteins), (vi) Regulatory and signaling modules (regulatory proteins: LysR and Crp/Fnr family transcriptional regulators, YhbH, HU-beta - signalling modules: adenylate cyclase, diguanylate cyclase/phosphodiesterase, NahK, KaiC) and, (vii) nucleic acid maintenance (DNA-binding protein HU-beta, CRISPR-Cas effectors (Cas9/Csx12), cyclic nucleotide phosphodiesterase, and ribonuclease Z-like proteins). Only 3% of the detected proteome was differentially expressed between parental and Δ*int* non/slow-growers, indicating that pP36 acted as a molecular switch, selectively reprogramming key pathways rather than inducing a global metabolic shift. This targeted regulation was consistent the action of a regulatory element. Strikingly, the mRNA of 75% of these enriched proteins contained predicted CsrA-binding motifs (5′-ANGGA-3′) (**Fig. 5d**), pointing to a post-transcriptional mechanism underlying pP36’s effects. CsrA, a conserved global post-transcriptional regulator, controls *L. pneumophila*’s transition between replicative and transmissive states, modulating virulence, motility, and growth rate, by binding to target transcripts ^56–72^. Among pP36’s cargo genes, *lpp0168* encodes a protein with a minimum of 31% identity and 51% similarity to CsrA and known homologous proteins (**Fig. 5e; Extended Data Fig. 6**) which adopts a CsrA-like fold, including the β-sheet architecture and RNA-binding dimerization interface (**Fig. 5f-g; Extended Data Fig. 6**). In biofilms, a Δ*lpp0168* mutant phenocopied Δ*int* and Δ*oriV*, with a 10-fold reduction in nongrowers (**Fig. 3; Extended Data Fig. 4a**) and increased growth resumption frequency (**Fig. 4**) on solid surfaces, implicating *lpp0168* in pP36-mediated sessile *L. pneumophila* growth rate heterogeneity. These findings position pP36 dynamics as a master regulator of non/slow-grower identity, maintaining a growth-arrested subpopulation through Lpp0168-dependent control.

**Figure 5:**
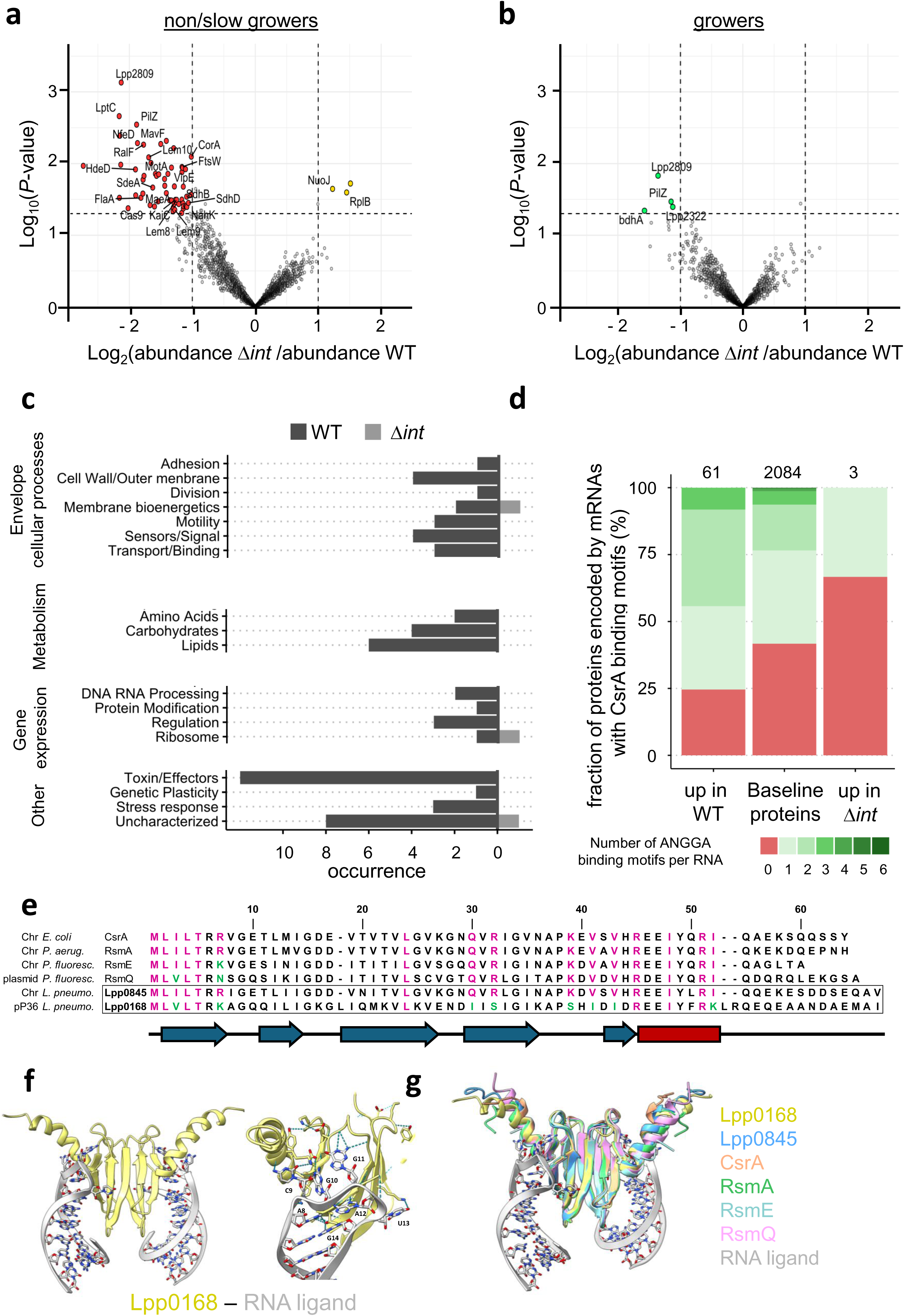
pP36 ICE dynamics shapes the molecular atlas of non/slow-growing *L. pneumophila*. *L. pneumophila* wild-type (WT) and the isogenic Δ*int* mutant, both producing the Timer reporter, were cultured to stationary phase, diluted in AYE broth, and incubated at 30°C for 48 h to form biofilms. Subpopulations were then sorted by FACS based on their Timer green/red fluorescence ratio, distinguishing growers from non/slow-growers. (**a-b**) Comparative proteomic analysis identified differentially produced proteins between WT and Δ*int* strains in both subpopulations. Protein abundance in each subpopulation is depicted as volcano plot. Data represent three biological replicates. (**c**) Differentially expressed proteins in WT and Δ*int* non/slow-growing subpopulations, categorized by functional category group. Black bars represent proteins enriched in WT, while grey bars indicate proteins enriched in the Δ*int* mutant. (**d**) Predicted CsrA-binding motifs (5′-ANGGA-3′) were mapped within ±100 nucleotides of the start codon in the coding sequences of proteins overexpressed in *L. pneumophila* WT or Δ*int* mutant non/slow-growers, relative to baseline protein levels. (**e**) Multiple sequence alignment of CsrA/Rsm family proteins. Alignment of Lpp0168 (a CsrA-like protein encoded on plasmid pP36) with the chromosomal *L. pneumophila* CsrA (Lpp0845), E. coli CsrA, and homologs RsmA (*P. aeruginosa*), RsmE (chromosome-encoded in *P. fluorescens*) and RsmG (plasmid-encoded in *P. fluorescens*). Pink highlights indicate conserved amino acids critical for RNA binding, while green highlights denote substitutions at these functionally important positions. (**f**) AlphaFold3-predicted structures of Lpp0168 (pP36-encoded CsrA-like protein) bound to target 20-nt RNA ligand. Inset: Close-up of key amino acids predicted to interact with mRNA nucleotides (A8, C9, G10, G11, A12 and U13), with hydrogen bonds depicted. (**g**) Structural alignment of dimers of Lpp0168 (yellow), Lpp0845 (blue), CsrA (orange), RsmA (green), RsmE (PBD: 2JPP; cyan) and RsmQ (pink) bound to 20-nt RNA ligand.

### pP36 ICE mediates horizontal transfer of antibiotic persistence mechanisms

The integrative model suggested that a heterogenous pP36 dynamics, cycling between integrated, excised, and replicating states, generate persister-like cells by imposing growth penalties, thereby enhancing population survival during fluoroquinolone treatment, where growing cells are preferentially killed (**Fig. 1c-d; Fig. SI 10, 11**). A classical method to assess antibiotic persistence is the biphasic killing curve, in which an initial rapid killing phase is followed by a plateau reflecting the survival of a tolerant subpopulation ^5^. We exposed *L. pneumophila* biofilms to ofloxacin (100 × MIC), a fluoroquinolone clinically relevant for Legionnaires’ disease treatment and previously used to detect *L. pneumophila* antibiotic persisters ^18,49^. Both parental and mutant strains exhibited biphasic killing kinetics indicating the presence of persister cells that tolerate ofloxacin. However, mutants with disrupted pP36 dynamics (Δ*int*, Δ*oriV*) or lacking the ICE-encoded CsrA-like regulator Lpp0168 (Δ*lpp0168*) displayed a 10-fold reduction in persister levels, consistent with the critical role for pP36 in nongrowers formation and identity (**Fig. 6a, left panel**). pP36 encodes all the machinery required to form a circular DNA intermediate upon excision, which can then be transferred and integrated into the chromosome of a recipient bacterium via conjugation. To determine whether pP36 could horizontally transfer persistence traits, we conjugated a kanamycin-marked pP36 from *L. pneumophila* Paris into the *L. pneumophila* clinical isolate CNR-L_21163689<u>401</u> (hereafter termed 401) lacking pP36 but possessing its integration site. Successful conjugation yielded Timer/*L. pneumophila* 401::pP36, as confirmed by qPCR showing chromosomal integration (decreased empty insertion sites) and active excision (circular pP36 forms) in biofilms (**Extended data Fig. 7a**). Comparative flow cytometry (**Fig. 6b**) and microscopy analyses (**Extended data Fig. 7b-c**) of biofilms revealed that pP36 acquisition by *L. pneumophila* 401 increased the nongrowing subpopulation size. This was accompanied with a reduced growth resumption frequency (**Fig. 6c**), and enhanced antibiotic persistence (**Fig. 6a, right panel**). qPCR analysis of approximately 10⁶ FACS-sorted cells confirmed that growth-arrested *L. pneumophila* 401::pP36 exhibited higher pP36 activity (**Fig. 6d**), mirroring dynamics in the original strain. Collectively, our findings suggest that, once acquired through horizontal gene transfer, pP36 interferes with the intrinsic phenotypic noise of the host bacterium and promotes the production of nongrowing cells and favor antibiotic persistence.

**Figure 6:**
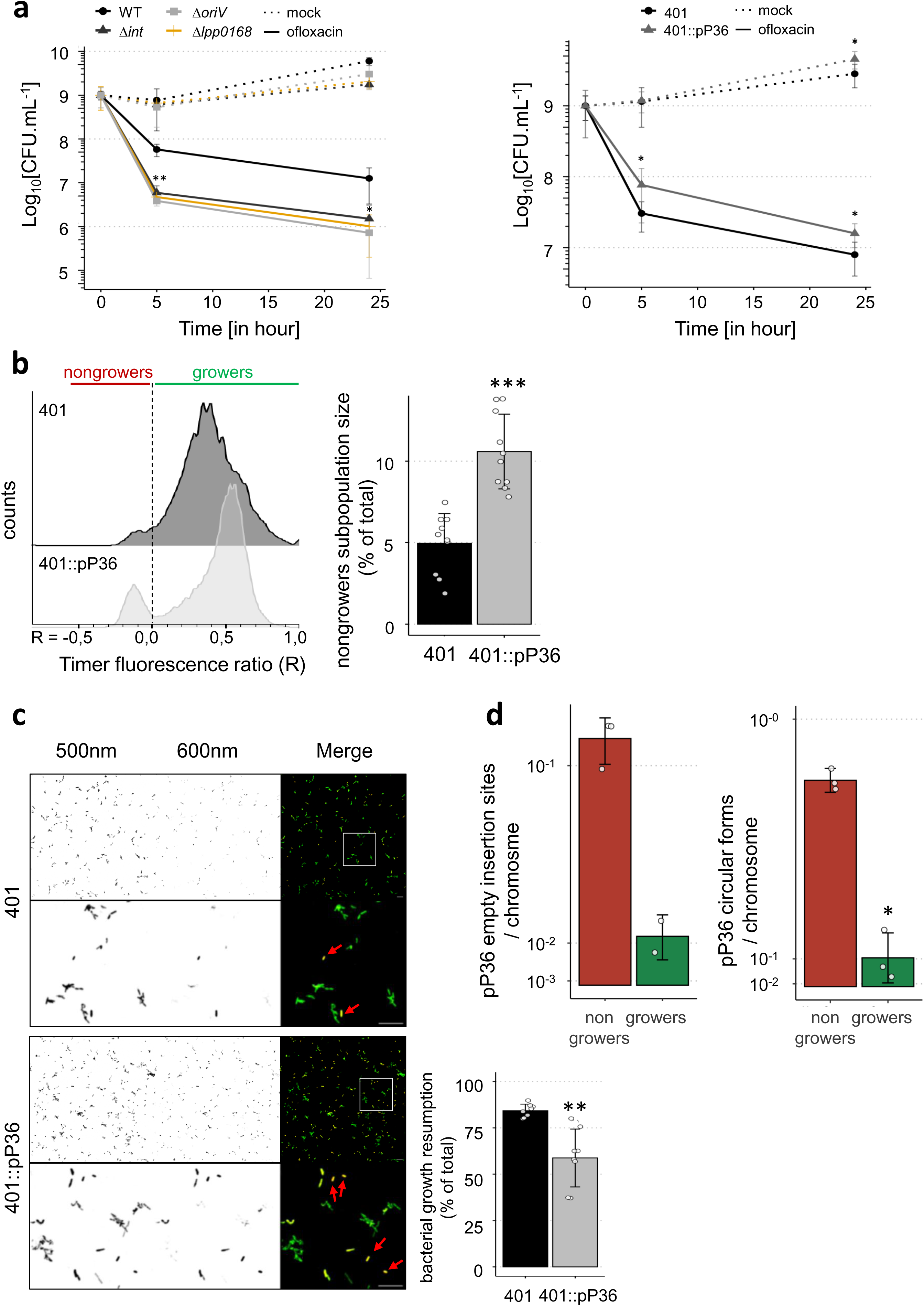
pP36 ICE mediates horizontal transfer of antibiotic persistence mechanisms. (**a**) Persister assay. Biofilms formed for 24 h at 30 °C by Timer-producing *L. pneumophila* strains were treated for the time indicated with ofloxacin (30 μg mL^−1^; full lines), or remained untreated (dashed lines). At defined time points, bacteria were washed, serial diluted and plated to quantify CFU. Left panel: *Legionella pneumophila* Paris wild-type and the isogenic mutant Δ*int*, Δ*oriV* and Δ*lpp0168*. Right panel: *L. pneumophila* strain 401 and *L. pneumophila* 401::pP36. Data represent the mean ± SD of three biological replicates (two-tailed Student’s t test; \*\*\**P* < 0.001, ** *P* < 0.01, \**P* < 0.05). (**b**) Biofilms formed for 24 h at 30 °C by Timer-producing *L. pneumophila* strain 401 and *L. pneumophila* 401::pP36 were homogenized and analysed by flow cytometry. The Timer fluorescence (500nm and 600nm) was recorded for individual bacterium. Left panel: frequency graph showing the distribution of the Timer fluorescence ratio (Log_10_[500 nm/600 nm]). Right panel: quantification of the subpopulation size in nongrowers. Data represent the mean ± SD of eight biological replicates (two-tailed Student’s t test; \*\*\**P* < 0.001, ** *P* < 0.01, \**P* < 0.05). (**c**) Stationary phase Timer-producing *L. pneumophila* strain 401 and *L. pneumophila* 401::pP36 were immobilized in AYE/0.5% agarose, and let form microcolony for 48 h. Fluorescence micrographs of microcolonies (scale bars, 20 μm). Magnification are showed. Red arrow: Nongrowing cells. Green arrow: Proliferating cells forming microcolonies. Quantification of bacterial growth resumption is displayed. Data represent the mean ± SD of four biological replicates (two-tailed Student’s t test; \*\*\**P* < 0.001, ** *P* < 0.01, \**P* < 0.05). (**d**) qPCR analysis of bacterial subpopulations from 24 h biofilms grown at 30°C. *L. pneumophila* 401::pP36 were sorted based on Timer fluorescence properties, genomic DNA extracted and pP36 empty insertion site and circular forms quantified by qPCR. Data represent the mean ± SD of three biological replicates (two-tailed Student’s t test; \*\*\**P* < 0.001, ** *P* < 0.01, \**P* < 0.05).

## Discussion

The antibiotic persistence, where a bacterial subpopulation survives treatment, poses a critical challenge to global health. While persistence is often attributed to stochastic phenotypic heterogeneity, the role of reversible genetic heterogeneity, such as that mediated by ICEs, remains unexplored. Here, to address this fundamental and unresolved question, we used a combination of stochastic simulations of a computational model and high-throughput single-cell measurements. Focusing on the *L. pneumophila* ICE pP36, we demonstrate that reversible genome structural variations: (i) drive the bacterium growth rate variation during biofilm formation, (ii) promote the coexistence of non-replicating antibiotic persisters alongside actively growing bacteria, (iii) define the molecular landscape of non/slow-replicating individuals and (iv) disseminate phenotypic heterogeneity and persistence traits.

*L. pneumophila* has recently emerged as a model organism for studying the molecular mechanisms underlying pathogen phenotypic heterogeneity and persister formation ^18,48,49^. As the causative agent of Legionnaires’ disease, a severe pneumonia that can relapse even after appropriate antibiotic treatment ^38,73,74^, this bacterium poses a clinical challenge. Isolates from recovered patients frequently lack genetically detectable antimicrobial resistance ^73,74^, suggesting that non-heritable mechanisms, such as persistence, contribute to treatment failure. Identifying the genetic determinants of antibiotic persistence is critical but complicated by the pleiotropic nature of the phenomenon. Our findings suggest that the presence of specific ICEs, such as pP36, could serve as a diagnostic marker for bacterial strains with an elevated risk of persister formation.

In our computational simulation framework, we explored how stochastic excision, integration, and replication of integrative and conjugative elements (ICEs) influence population-level heterogeneity and the emergence of persister-like antibiotic-tolerant individuals in *L. pneumophila*. Our experimental findings corroborate the model, demonstrating that ICE dynamics induce growth rate variability and recapitulate the biphasic killing kinetics characteristic of bacterial persistence. While our simulations predicted that immobilizing ICE dynamics would abolish biphasic killing kinetics by eliminating ICE-driven growth heterogeneity, experimental disruption of pP36 excision (*i.e.*, Δ*int*) only reduced persister levels, reflecting the multifactorial nature of persistence. This partial effect suggests that ICEs like pP36 amplify, but do not exclusively control, persister formation, and other pathways contribute to residual ofloxacin tolerance.

pP36 maintain a subpopulation of growth-arrested *L. pneumophila* through a CsrA-like (Lpp0168)-dependent control. We previously characterized the *Legionella* quorum sensing system (Lqs) as a contributor to persister formation during biofilm development ^49^. In *L. pneumophila*, chromosomal CsrA directly represses translation of the quorum-sensing regulator LqsR, while the small RNAs RsmY/Z sequester CsrA, thereby modulating this repression ^66^. The pP36-encoded CsrA-like regulator could disrupt this balance in a gene-dosage-dependent manner by competing for RNA targets, titrating RsmY/Z, or partially antagonizing chromosomal CsrA. Such interference may reshape Lqs signalling dynamics, thereby linking ICE activity to host growth and persistence programs, and providing yet another example of the remarkable versatility of CsrA functions in bacterial cells.

A key finding in this study is the ability for pP36 to disseminate growth-rate heterogeneity and persistence traits across bacterial populations through horizontal transfer. However, when pP36 is mobilized into a heterologous genetic background, its impact on growth-rate heterogeneity and persister formation is attenuated compared to its native host. This attenuation likely reflects the necessity for the ICE-encoded functions to interface with unfamiliar regulatory circuits. Hence, the adaptive value of mobile genetic elements derives not only from their dynamics and genetic cargo but also from their ability to embed and coevolve within host regulatory architectures.

During biofilm formation, the pP36 actively prevents growth resumption in a subpopulation of *L. pneumophila*, likely through its regulation of stress response, energy metabolism, cell envelope integrity, and nucleotide and lipid metabolism. By maintaining cells in a nongrowing state, pP36 ensures a bacterial reservoir poised for survival under stress. Concurrently, this mechanism may reduce the risk of losing pP36 in its vulnerable circular form. When pP36 dynamics are disrupted, cells resume division prematurely. The few remaining individuals with reduced growth rate exhibit a distinct, compromised proteomic profile, lacking key adaptations for stress resilience, metabolic adaptability, motility, and virulence. This suggests that pP36 is not only a gatekeeper of persister-like cell formation but also a curator of persister-like cell robustness.

Our qPCR data capture only a single time point, meaning that individual cells with excised pP36 may later reintegrate the element, while excision simultaneously occurs in other cells within the subpopulation. This continuous cycling of pP36 between excised and integrated states collectively maintains its dynamic influence across the growth-arrested bacterial population. Future work will dissect the real-time dynamics of pP36 excision and reintegration, and determine how this stochastic switching contributes to the stability and revival of nongrowing subpopulations. The mechanisms underlying the heterogeneous induction of pP36 excision remain unclear. Transcriptional feedback loops may amplify minor stochastic differences in the expression of the pP36 integrase/excisionase, allowing the ICE to adopt two distinct stable states, integrated or circularized, under identical conditions. Such bistable gene expression has been observed in bacterial nanomachines ^75^ and the mobile genetic elements ICE*clc* and *Tn916* ^76,77^.

To our knowledge, this study provides the first evidence directly linking genome structural dynamics to both phenotypic noise and the horizontal dissemination of antibiotic persistence traits. MGE’s phenotypic noise is not exclusive to ICEs. A recent study has shown that synthetic multicopy plasmids exhibit substantial copy number heterogeneity, leading to noisy expression of plasmid-encoded genes, including resistance cassettes, and resulting in transient antibiotic resistance ^78^. Together, these observations suggest that coupling heterogeneous MGE dynamics to bacterial phenotypic variations may represent a conserved mutualistic strategy, benefiting both the mobile genetic elements and their bacterial hosts.

Decades of antibiotic use have enabled unprecedented control of infectious diseases and fuelled advances across modern medicine that depend on effective infection management. Yet, antibiotic therapeutic failures remain a pressing global health threat. While ICEs have traditionally been linked to the expression and horizontal transfer of antibiotic resistance genes, our work reveals that they also drive pathogens’ phenotypic variation. At the single-cell level, dynamic ICE-mediated reversible genome structural variations provide individual bacteria with adaptive advantages under fluctuating conditions. At the population level, the horizontal dissemination of these traits fosters collective resilience, enhancing the community’s capacity to withstand environmental stressors and antimicrobial challenges.

## Methods

### Stochastic multiscale modeling framework

We developed *msevol*, a stochastic multiscale modeling framework built on a nested, graph-based architecture that links genetic, cellular, and ecological scales (https://github.com/rasigadelab/msevol/wiki). In this representation, cells and mobile genetic elements are modeled as discrete entities connected through dynamic inclusion networks that govern processes such as excision and intracellular replication of ICEs, fitness costs, and selection under antibiotic pressure. A complete description of the model structure, assumptions, and parameterization is provided in the **Supplementary Information** file. The model was implemented in C++, and all simulations, figures, and data analyses were performed in R. The source code of the simulator and the simulation scripts are publicly available on GitHub (https://github.com/rasigadelab/msevol2_funmge; https://github.com/rasigadelab/Rmse2FunMGE).

### Bacterial strains, cell lines and culture conditions

Bacterial strains used in this study are listed in the **Extended Data Table 3**. *L. pneumophila* strains are derived from *L. pneumophila* Paris Outbreak isolate CIP107629 (CNR Lyon) ^79^ and *Legionella pneumophila* clinical isolate #021163689401 (CNR Lyon). Bacteria were cultured on charcoal yeast extract (BCYE) agar plates, buffered with N-(2-acetamido)-2-aminoethane sulfonic acid (ACES) at 37°C and supplemented with Chloramphenicol (Cam) 10 µg mL^-1^ , Kanamycin (Kan) 20 µg mL^-1^ or isopropyl β-d-1-thiogalactopyranoside (IPTG) 1 mM, when required. After 72 h, bacteria were sub-cultured on fresh CYE plates and kept at 37°C. To prepare routine working liquid cultures, ACES-buffered yeast extract (AYE) medium, supplemented with appropriate antibiotics, was inoculated with bacterial colonies from a fresh agar plate to an initial OD₆₀₀_nm_ of 0.1 and incubated under fully aerated conditions at 37°C with shaking for 21–22 hours to reach post-exponential phase (OD₆₀₀_nm_ of ± 4.5 – 5.5, motile coccobacilli). *Escherichia coli* DH5α were cultured on LB plates or liquid medium supplemented with the appropriate antibiotics. *E. coli* DH5α were transformed by heat shock using 100 ng of plasmid DNA. *L. pneumophila* strains were transformed by electroporation (2.4 kV, 100 Ω, 25 µF) with 2 µg of plasmid DNA.

### Identification of genomic islands in *L. pneumophila* strain Paris

The identification and mapping of genomic islands in *L. pneumophila* strain Paris was conducted by literature search ^40,43,80–84^ and refined *in silico* through sequence analysis. Whenever an integrase was detected, the nearest tRNA-encoding sequences (preferred insertion sites for genomic islands) were analyzed. Since integrase-mediated integration necessarily generates inverted repeat sequences, these tRNA regions were aligned within 150 kb upstream and downstream of the integrase to identify potential repeats. When such repeats were found, the island boundaries were precisely mapped.

### Identification of the putative *oriV* of the ICE pP36

The putative origin of vegetative replication (*oriV*) of the ICE pP36 was mapped through bioinformatic and sequence analysis as follow: (i) The pP36-encoded gene *lpp0166*, initially annotated as a hypothetical protein, was identified as putative replicase (Phyre2.0 ^85^). (ii) The 500-bp regions immediately upstream *lpp0166* contains three predicted DnaA boxes, systematically flanked by AT-rich regions, and four AT-rich regions harboring direct and inverted repeats of an 8-bp perfect sequence. Such arrangement mirrors the structural organization of iteron-containing replicons.

### Quantitative PCR (qPCR)

Quantification of the empty chromosomal *att* site, the circular form of pP36, and LPPGI-A was performed using real-time qPCR with SsoFast EvaGreen Supermix (Bio-Rad, #1725202) on a StepOnePlus Real-Time PCR System (Applied Biosystems, #15331295). Cycling parameters were as follows: 95 °C for 30 s; 40 cycles of 95 °C for 5 s and 60 °C for 5 s; melt curve from 65 °C to 95 °C with 0.5 °C increments every 5 s. Data were analyzed with StepOne Software using the ΔΔCT method, with *gyrB* and *ribD* as reference genes. The unoccupied chromosomal *att* locus was amplified to quantify empty integration sites, and the circularized *att* junction was amplified to quantify excised ICE copies. Primer sequences are listed in in the **Extended Data Table 4**.

### Biofilm and microcolony formation

For biofilm formation, liquid cultures were diluted in fresh AYE/Cam at an OD_600nm_ of 0.1, placed in a multi-well plate or an ibidi imaging chamber, and incubated for the indicated temperature and time, while avoiding mechanical disturbance as in ^49^. The formation and analysis of microcolonies is detailed in ^49,55^. Briefly, cultured bacteria were embedded in jellified AYE (AYE/Cam + 0.5% agarose) at an OD_600nm_ of 0.1. Jellified AYE/Cam was obtained by boiling 10 mL AYE and 100 μg of Ultrapure AgaroseTM (Invitrogen) followed by the addition of 10 mL AYE/Cam. Upon addition of the bacteria, the suspension was thoroughly mixed and immediately loaded into an ibidi μ-Slides 8 Wells or a 96-well plate, let solidify and incubated 48 h at 30°C.

### Confocal microscopy and image analysis

Images acquisition was carried out using the confocal microscope AxioObserver Z1 LSP800 (Zeiss; SFR Biosciences, Lyon). Biofilm and microcolony were imaged in 3 dimensions (z-stack of 14 to 15 slices) at 60 × (optical section size: 230 nm) or 40 × (optical section size: 360 nm) magnification, respectively. Various fields per well and time point were imaged. The Timer’s spectral properties were recorded for each individual bacterium using the following setups: LSM800: (green, ex. 488 nm, em. 515–545 nm; red, ex. 561 nm, em. 600–620 nm.

Biofilm images analysis were analyzed using a CellProfiler-based pipeline. One optical section over two was processed. Bacteria were segmented using the *Omnipose* model *bact_fluor_omni* ^86^ before integration into CellProfiler 4.2.5 ^87^ and the Timer fluorescence calculated. For microcolony images, segmentation was performed using Minimum Cross-Entropy thresholding on projection images of a Cellprofiler pipeline. Object were dilated, merged and eroded to obtain final segmentation of microcolonies. Data were analyzed in R (RStudio) using *ggplot2* for graph generation. Image montages were prepared using Fiji (ImageJ) software version 2.14.0.

### Flow cytometry analysis

Biofilms were homogenized, centrifuged (1500 *g*, 10 min), resuspended in ice-cold PBS, fixed 4% paraformaldehyde (PFA) for 20 min and washed in PBS. Timer spectral parameters were subsequently recorded in a FACS-Fortessa II (BD Biosciences). A minimum of 10’000 bacteria per sample were acquired. The gating strategy was performed as described in ^18,49^. Data processing was realized with the software FlowJo (V10.8.1). Timer fluorescence ratio was calculated for each detected bacterium as following: Log_10_[(Ex 488 nm, Em 515–545 nm)/(Ex 561 nm, Em 600–620 nm)].

### Cell sorting

Biofilms were homogenized, centrifuged (5000 rpm, 15 min) and resuspended in ice-cold PBS. Cells were sorted by Timer color ratio on a FACSAria III (BD Biosciences) using scatter and fluorescence parameters (green: excitation 488 nm, emission 515–545 nm; red: excitation 561 nm, emission 600–620 nm), a 70 µm nozzle, four-way purity mode, and ± 90 % sorting efficiency. Sorted subsets were systematically reanalyzed.

### Genome edition

Deletions of Δ*lpp0194*, Δ*oriV* and Δ*lpp0168* in *Legionella pneumophila* were generated by homologous recombination as described in ^88^. Briefly, approximately 2 kb upstream (P1-P2) and downstream (P3-P4) the gene of interest were amplified by PCR and fused to a *kanamycin*/*mazF* resistance–suicide cassette by double-joint PCR (P1-P4). The construct replaced the target gene following natural transformation, and mutants were selected on kanamycin and confirmed by PCR. PCR amplifications were performed by using the PrimeStar Max DNA polymerase Premix, following the manufacturer’s instructions (Takara). Primer sequences are listed in in the **Extended Data Table 4**.

### Comparative proteomic analysis

For sample preparation, biofilms were homogenized, centrifuged (5000 rpm, 15 min) and resuspended in ice-cold PBS. Bacterial suspension and sorted bacteria were kept in ice-cold PBS supplemented with 180 μM Cam to prevent protein degradation and synthesis. Approximately 1 × 10⁶ sorted bacteria per subset were collected (three biological replicates). Sorted bacteria were prepared and LC-MS analyzed as described recently ^89^ with the following modifications. Approximately 50 ng of peptides were subjected to LC–MS/MS analysis using a timsTOF Ultra Mass Spectrometer (Bruker) fitted with a Vanquish Neo HPLC (Thermo Fisher Scientific) and the column heater was set to 60°C. Peptides were resolved using a RP-HPLC column (100μm × 30cm) packed in-house with C18 resin (ReproSil Saphir 100 C18, 1.5 μm resin; Dr. Maisch GmbH) at a flow rate of 0.4 μL min-1. A linear gradient of buffer B (80% acetonitrile, 0.1% formic acid in water) ranging from 2% to 25% over 25 minutes and from 25% to 35% over 5 minutes was used for peptide separation. Buffer A was 0.1% formic acid in water. The mass spectrometer was operated in DIA-PASEF mode with the following source settings: 500V End Plate Offset, 4500V Capillary Voltage, 0.4bar Nebulier Pressure, 3 l/min Dry Gas flow at 180°C. The dia-PASEF scheme can be seen in supplementary table X. The collision energy was ramped between 20 eV at 0.6 V -s/cm² and 59 eV at 1.6 V -s/cm². DIANN (v1.8.1) was used to search all raw MS data against a FASTA database containing all predicted proteins (4377 sequences, downloaded 04/06/2024 from NCBI) from *Legionella pneumophila str. Paris*. Taxonomy ID: 297246.

### Sequence alignment and identity analysis

Protein sequence identity and similarity were calculated using the EMBOSS Needle Pairwise Alignment tool (PSA; EMBL-EBI ^90^), while multiple sequence alignments were performed with MUSCLE (EMBL-EBI ^90^). Amino acid residues involved in RNA recognition, as previously identified ^61,91^, were annotated in the alignments: conserved residues were highlighted in magenta, and variable or substituted residues were highlighted in green.

### Structural modeling and analysis

Structural predictions were performed using the AlphaFold 3 server ^92^. The resulting models were visualized and analyzed using ChimeraX v1.4 ^93^. Structural comparisons were conducted using the Needleman-Wunsch algorithm with the BLOSUM-62 substitution matrix for protein matchmaking. For comparative analysis, the structures of RsmA from *Pseudomonas aeruginosa*, RsmE from *Pseudomonas fluorescens*, RsmQ from a plasmid of *Pseudomonas fluorescens*, and CsrA from *Escherichia coli* were also modeled. Complexes including the 20-nucleotide RNA sequence (GGGCUUCACGGAUGAAGCCC) were inferred from ^61^. The structure used for RsmE when bound to the 20-nt RNA used is the published structure (PBD: 2JPP) ^61^.

### Persister assay

*L. pneumophila* were grown to stationary phase, diluted in AYE broth, and allowed to form biofilms for 24 h at 30 °C prior treatment with ofloxacin 30 μg.mL^−1^ (100 × MIC). At given time points, bacteria were collected and washed three times. For each wash 1 mL PBS was used and 950 μL of the supernatant discarded. The bacterial suspensions were subsequently serially diluted 1:10 and plated to quantify CFUs.

### Conjugation

First, pP36 was tagged using a kanamycin resistance cassette, integrated by homologous recombination ^88^ using the primers oKH235 and oKH312. To minimize potential disruption of plasmid functions, the cassette was inserted between the convergent genes *lpp0187* and *lpp0188*. Successful integration was confirmed by PCR amplification. The integrative and conjugative element (ICE) pP36, carrying a kanamycin resistance cassette, was transferred from *L. pneumophila* Paris to the Timer-producing clinical isolate *L. pneumophila* 401 by bacterial conjugation. Briefly, 100 µL of donor and recipient liquid cultures were mixed, centrifuged for 5 minutes at 5,000 rpm, and the supernatant discarded. The bacterial pellets were then resuspended in 50 µL of AYE medium, spotted onto CYE agar, allowed to dry for 30 min, and incubated at 30°C for 24 h. The bacterial spot was then resuspended in 500 µL of AYE broth and plated onto selective media (CYE plates supplemented with 30 µg.mL^-1^ Cam and 50 µg.mL^-1^ kan) to select for transconjugants. The presence of plasmid pP36 in the recipient strain was subsequently confirmed by PCR. The resulting strain was designated Timer/*L. pneumophila* 401::pP36. Primer sequences are listed in in the **Extended Data Table 4**.

### Quantification and statistical analysis

Statistical differences were determined using a two-tailed Student *t* test on the means of at least three independent experiments. Probability (*P*) values of less than 0.05, 0.01, and 0.001 were used to show statistically significant differences and are represented with *, **, or ***, respectively. For the comparative proteomics, the summarized protein expression values were used for statistical testing of between condition differentially abundant proteins using MSstats R package v.4.7.3 (https://doi.org/10.1093/bioinformatics/btu305).

## Data availability

All mass spectrometry proteomics data associated with this manuscript have been deposited to the ProteomicsXchange consortium (dataset PXD074064) via MassIVE (https://massive.ucsd.edu) with the accession number MSV000100721. All other relevant data are included in this article and its Supplementary Information files, or from the corresponding author upon request.

## Acknowledgment

We are grateful to past and present members of the N.P. research laboratory for their insightful comments and stimulating discussions. We also thank the SFR Biosciences platforms ANIRA-Cytometry and LyMIC-PLATIM for providing access to flow cytometers, cell sorters, and microscopes. Research in the N.P. laboratory is supported by the Centre international de recherche en infectiologie (CNRS, Inserm, Université Claude Bernard Lyon 1), a CIRI seed-fund, a CIRI transversal research project fund, and the Agence nationale de la recherche (grants ANR-24-CE15-0280-01, ANR-24-CE44-6047-02, and ANR-22-CE20-0018-02). I.D. is supported by an E2M2 PhD fellowship. D.B. is a tenured researcher in the N.P. laboratory and a lecturer at Université Claude Bernard Lyon 1. N.P. is a tenured CNRS researcher. Development of *msevol* is supported by ANR under grant ResisTrack (ANR-20-CE35-0012), by the LABEX ECOFECT (ANR-11-LABX-0048) of Université de Lyon, within the program “Investissements d’Avenir” (ANR-11-IDEX-0007) operated by the French National Research Agency (ANR) and by the RESPOND program of the Université de Lyon (UDL), awarded to JPR. The funders had no role in study design, data collection and analysis, decision to publish, or preparation of the manuscript.

## Authors information

### Authors and primary affiliations

Iris Dadole, Romain Gory, Kevin T. Huguet, Benoît Morin, Didier Blaha, Nicolas Personnic: Team « Persistence and single-cell dynamics of respiratory pathogens », Centre International de Recherche en Infectiologie, INSERM U1111, Université Claude Bernard Lyon 1, CNRS, UMR 5308, École Normale Supérieure de Lyon, Université de Lyon, Lyon, France

Natacha Lenuzza, Jean-Philippe Rasigade: Team « Public Health, Epidemiology and Evolutionary Ecology of Infectious Diseases », Centre International de Recherche en Infectiologie, INSERM U1111, Université Claude Bernard Lyon 1, CNRS, UMR 5308, École Normale Supérieure de Lyon, Université de Lyon, Lyon, France

Laetitia Attaiech, Xavier Charpentier: Team “Horizontal gene transfer in bacterial pathogens,” Centre International de Recherche en Infectiologie, INSERM U1111, Université Claude Bernard Lyon 1, CNRS, UMR 5308, École Normale Supérieure de Lyon, Université de Lyon, 69100 Villeurbanne, France

Christophe Ginevra, Sophie Jarraud: Centre National de Reference des Légionelles, Institut des Agents Infectieux, Hospices Civils de Lyon, Lyon, France

Friederike Steurer, Alexander Schmidt: Proteomics Core Facility, Biozentrum, University of Basel, Basel, Switzerland

### Contributions

I.D., R.G., K.H., D.B., N.P. designed the experiments – I.D., R.G., K.H., B.M. and N.P. performed experiments and analysed the data – N.L. and J.P.R. developed the simulation – K.H, L.A and X.C. created bacterial strains – C.G. and S.J. identified and provided *L. pneumophila* clinical isolate – F.S. and A.S. performed the proteomic analysis – I.D. and N.P. wrote the article, which was reviewed and approved by all authors – N.P. supervised the work and acquired funds.

### Corresponding author

Correspondence to Nicolas Personnic

---

**Extended Data Figure 1:**
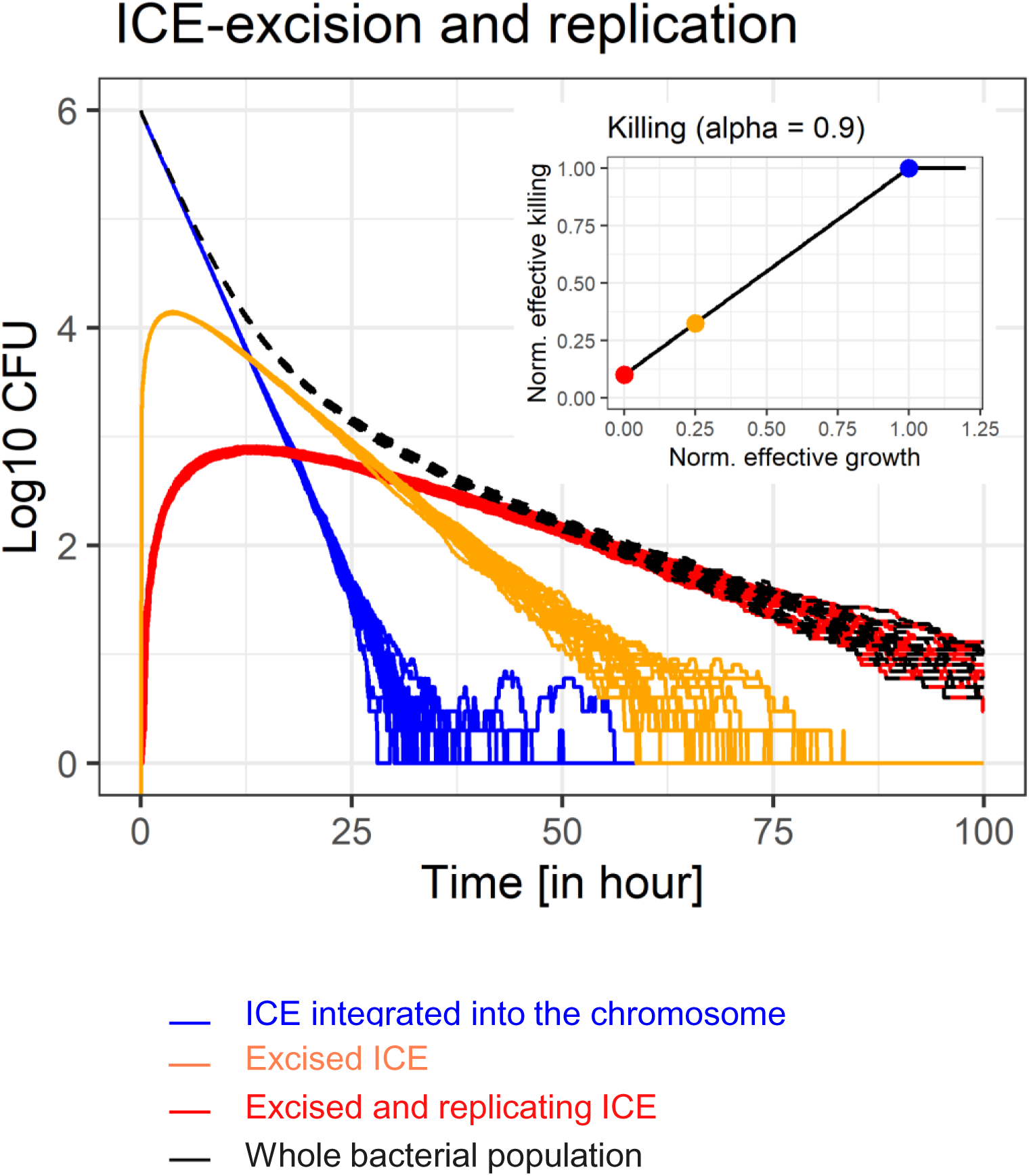
Stochastic ICE dynamics favors bacterial growth rate heterogeneity and antibiotic persistence. Linked to Figure 1. Simulated time–kill assay of an initially homogeneous bacterial population capable of diversifying from its initial genotype through ICE excision and replication. All cells initially carried an integrated ICE, allowing them to divide at their maximal intrinsic growth rate and, consequently, to experience maximal antibiotic killing. Over time, the population diversified through ICE excision and subsequent increases in free ICE copy number, which progressively reduced both growth rate and antibiotic-mediated killing. *N* = 20 simulations; black curves total population. Colors indicate the effective growth rates of subpopulations, reflecting differences in ICE copy number. Blue: one integrated, silent ICE, normal growth; orange: one excised ICE, reduced growth; red: two or more excised ICE copies, no growth. This example simulations were performed with an ICE fitness cost of −0.15 per hour (one excised copy reduces growth; two or more excised copies completely stop growth), a growth–killing coupling factor of 0.9, an excision rate of 0.01 per hour, a duplication rate of 0.0 per hour, and a loss rate of 0.001 per hour, while all other parameters were kept at their baseline values.

**Extended Data Figure 2:**
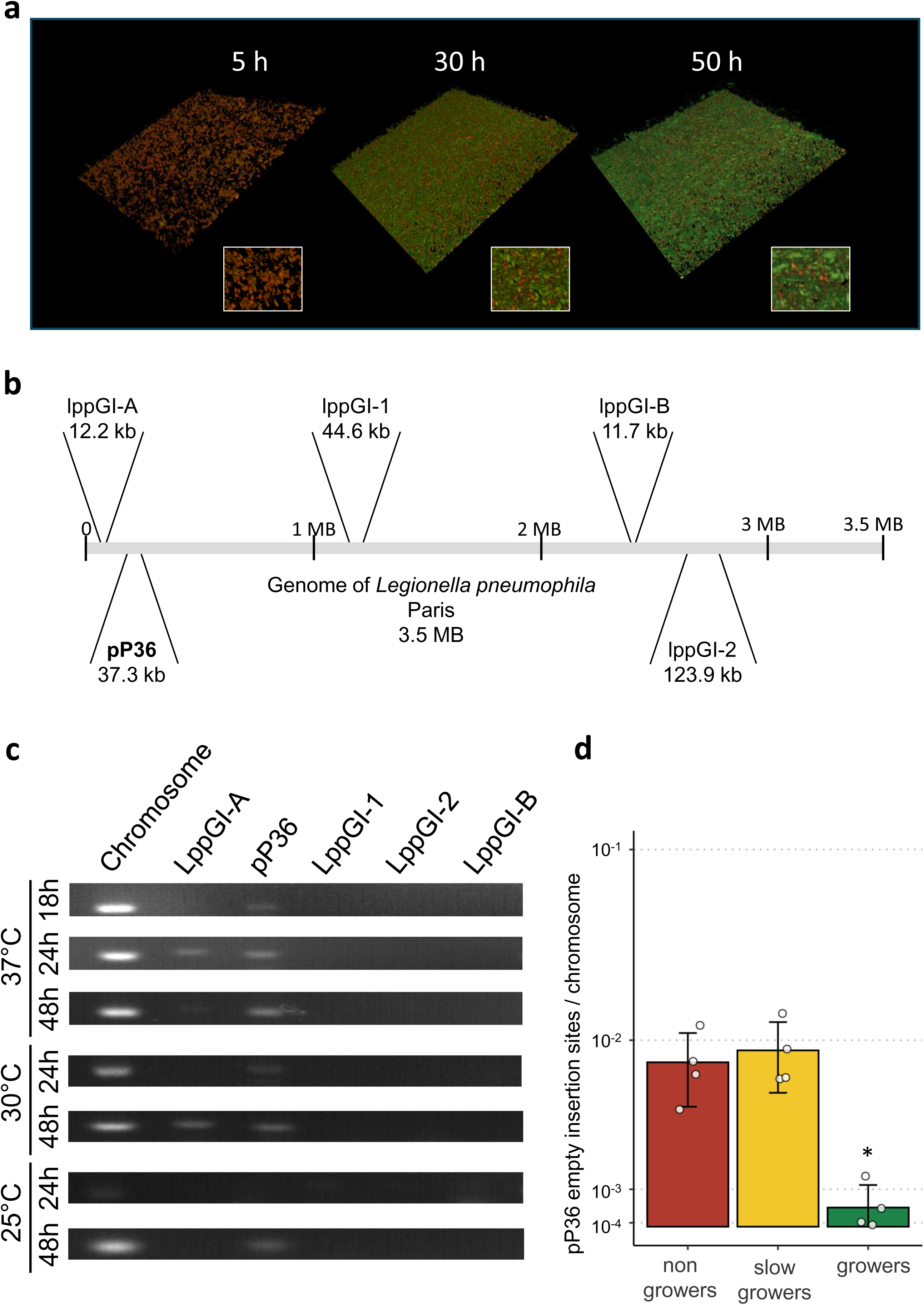
pP36 ICE dynamics is heterogeneous and bound to *L. pneumophila* growth status. Linked to Figure 2. *L. pneumophila* producing Timer were grown to stationary phase, diluted in AYE broth, and allowed to form biofilms at 30 °C. (**a**) Confocal microscopy. 3D reconstruction (x = 100 μm, y = 100 μm, z = 10 μm). Micrographs show overlays of the Timer fluorescence (500 nm and 600 nm); growing and nongrowing bacteria appear green or red/orange, respectively. (**b**) *Legionella pneumophila* Paris ICEs genetic organisation. (**c**) PCR-screening of five ICEs in circular form in biofilms grown at different temperatures (25°C, 30°C, and 37°C) and growth times (18 h, 24 h, and 48 h). (**d**) qPCR analysis of bacterial subpopulations from 48-hour biofilms grown at 30°C. *L. pneumophila* were sorted based on Timer fluorescence properties, genomic DNA extracted and pP36 empty insertion site quantified by qPCR. Data represent the mean ± SD of four biological replicates (two-tailed Student’s t test; \*\*\**P* < 0.001, ** *P* < 0.01, \**P* < 0.05).

**Extended Data Figure 3:**
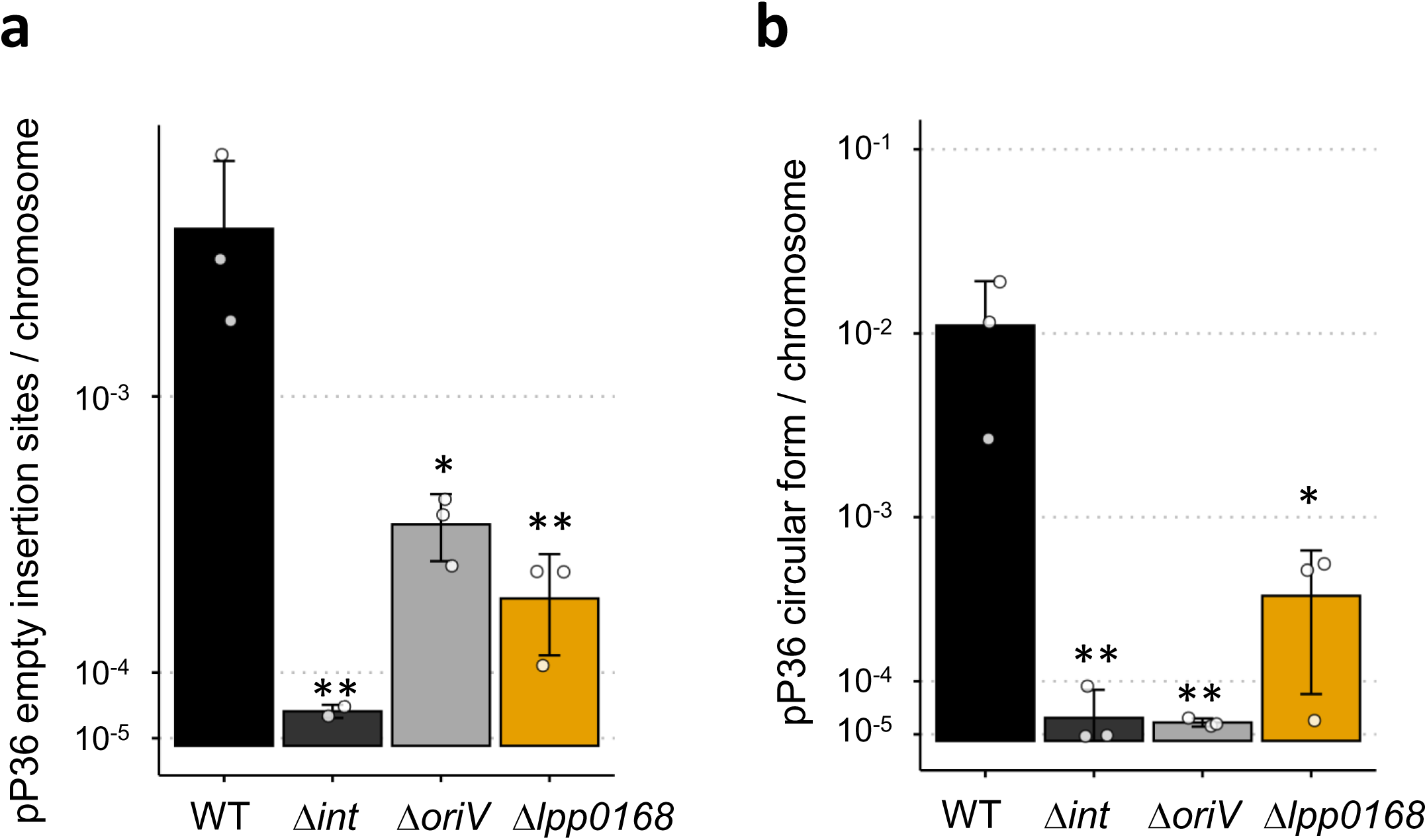
pP36 ICE dynamics. Linked to Figure 3. *L. pneumophila* wild-type (WT) and the isogenic mutants Δ*int*, Δ*oriV* and Δ*lpp0168*, producing Timer, were grown to stationary phase, diluted in AYE broth, allowed to form biofilms at 30 °C for 24 h. (**a**) qPCR analysis of pP36 empty sites. (**b**) qPCR analysis of pP36 circular forms. Data represent the mean ± SD of three biological replicates (two-tailed Student’s t test; \*\*\**P* < 0.001, ** *P* < 0.01, \**P* < 0.05).

**Extended Data Figure 4:**
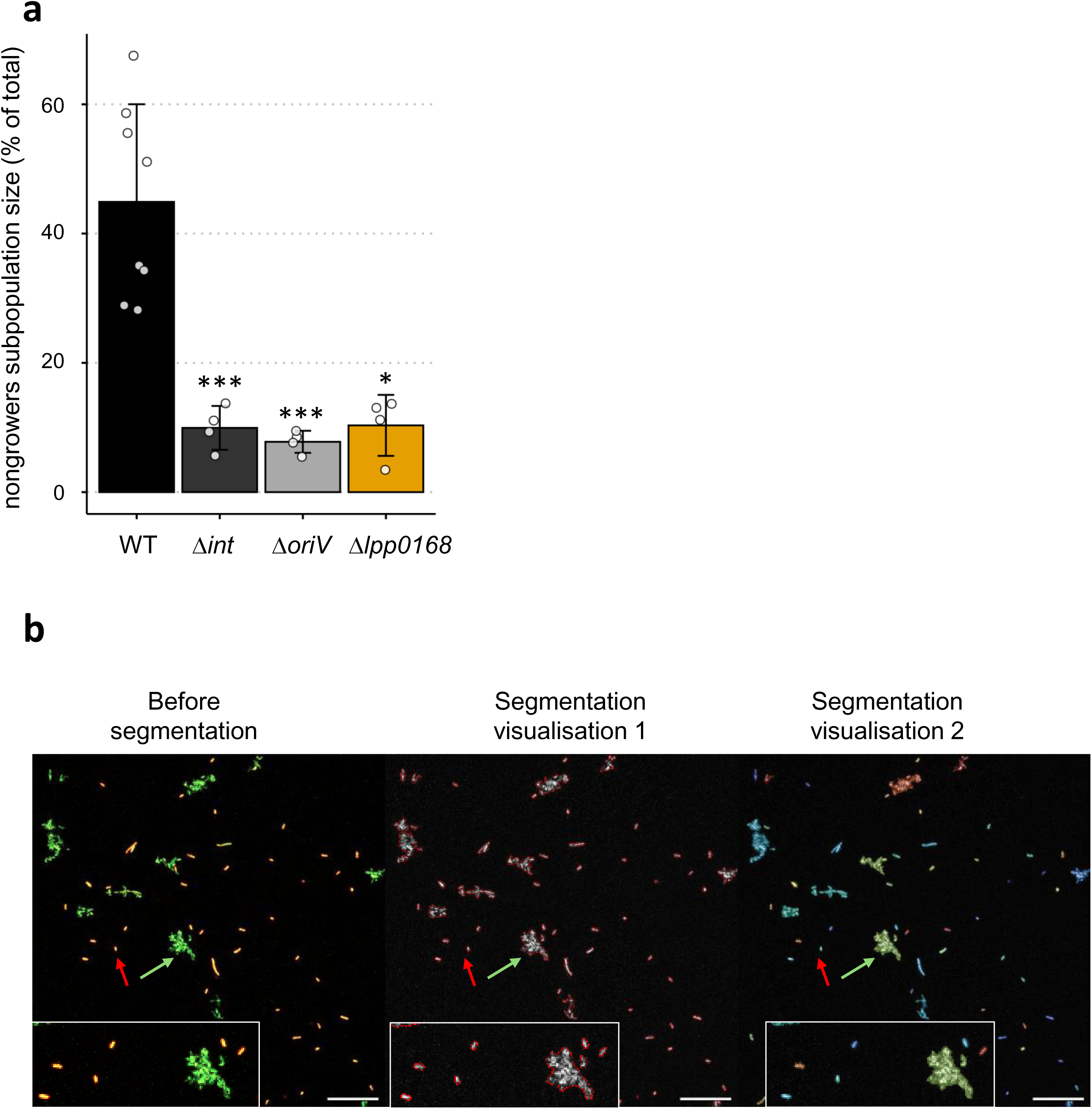
pP36 ICE dynamics orchestrates sessile *L. pneumophila* growth rate heterogeneity. Linked to Figure 3 & Figure 4. (**a**) *L. pneumophila* wild-type (WT) and the isogenic mutants Δ*int*, Δ*oriV* and Δ*lpp0168* producing Timer, were grown to stationary phase, diluted in AYE broth, and allowed to form biofilms at 30 °C for 24 h. Quantification of the subpopulation size in nongrowers by confocal microscopy. Data represent the mean ± SD of four biological replicates (two-tailed Student’s t test; \*\*\**P* < 0.001, ** *P* < 0.01, \**P* < 0.05). (**b**) *L. pneumophila* expressing Timer was cultured to stationary phase, diluted at OD 0.1 in AYE broth + agarose 0.5%, and incubated at 30°C for 48 hours to allow microcolony formation. Red arrow: nongrowing cells. Green arrow: proliferating cells forming microcolonies. Microcolony formation was analysed using Cellprofiller (particle analysis, segmentation and area measurement). Representative segmentations are shown for the subsequent quantification of nongrowers and microcolonies.

**Extended Data Figure 5:**
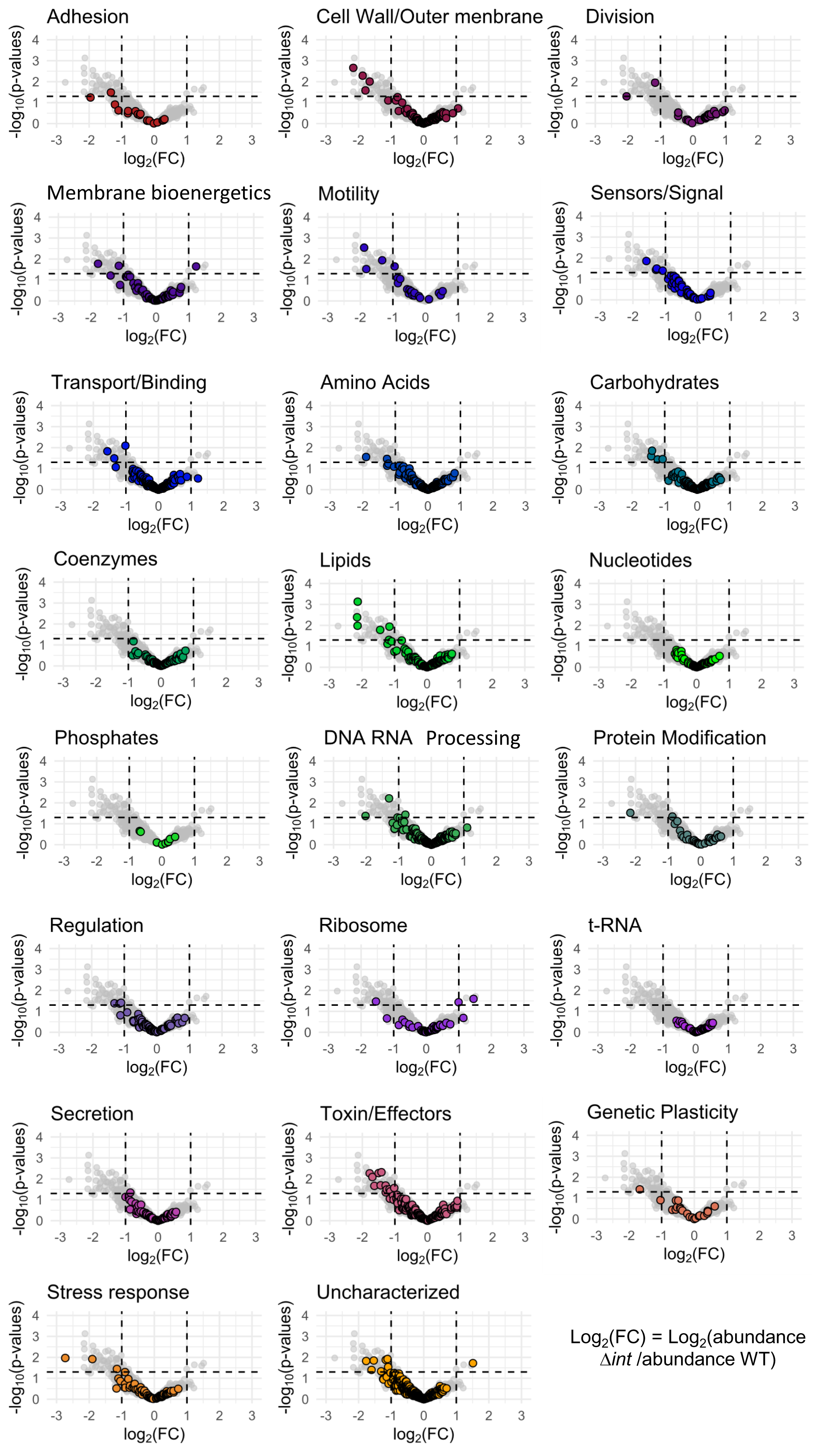
pP36 ICE dynamics shapes the molecular atlas of non/slow-growing *L. pneumophila*. Linked to Figure 5. *L. pneumophila* wild-type (WT) and the isogenic Δ*int* mutant, both producing the Timer reporter, were cultured to stationary phase, diluted in AYE broth, and incubated at 30°C for 48 h to form biofilms. Subpopulations were then sorted by FACS based on their Timer green/red fluorescence ratio, distinguishing growers from non/slow-growers. Comparative proteomic analysis identified differentially produced proteins between WT and Δ*int* non/slow-growing bacteria. Protein abundance is depicted as volcano plot categorized by functional category group.

**Extended Data Figure 6:**
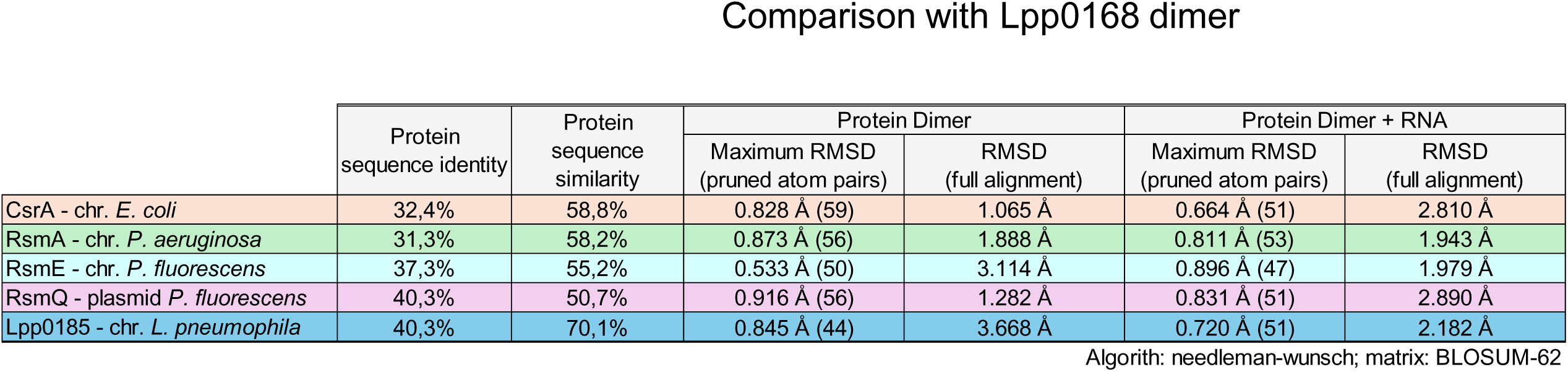
Structural and sequence comparison of Lpp0168 with homologous proteins. Linked to Figure 5. Comparison of the sequence and structural features of Lpp0168 (pP36-encoded CsrA-like protein) with CsrA (*E. coli*), and known homologous proteins RsmA (*P. aeruginosa*), RsmE (*P. fluorescens*), RsmQ (*P. fluorescence*) and Lpp0845 (*L. pneumophila*). chr, plasmid: corresponding gene encoded on the bacterial chromosome or on a plasmid, respectively. Protein sequence identity represents the percentage of identical amino-acid residues between each protein and Lpp0168, while protein sequence similarity reflects the percentage of similar residues, including conservative substitutions. The root-mean-square deviation (RMSD) values (in angstroms, Å) indicate how much two structures differ from each. For each tested condition (*i.e.,* dimeric structures, either alone or in complex with RNA (“Dimer” and “Dimer + RNA” columns, respectively), two RMSD values are reported. The “Maximum RMSD” focuses on the most similar subset of atoms, less affected by flexible regions. The numbers in parentheses indicate the number of pruned atom pairs considered in the RMSD calculation. The RMSD based on the full structural alignment calculated after superimposing all atoms and highly sensitive to local deviations notably in flexible regions. RMSD value < 1.0 Å, excellent agreement; 1.0 - 2.0 Å: small differences; 2.0–3.0 Å, moderate deviations; > 3.0 Å, significant structural divergence.

**Extended Data Figure 7:**
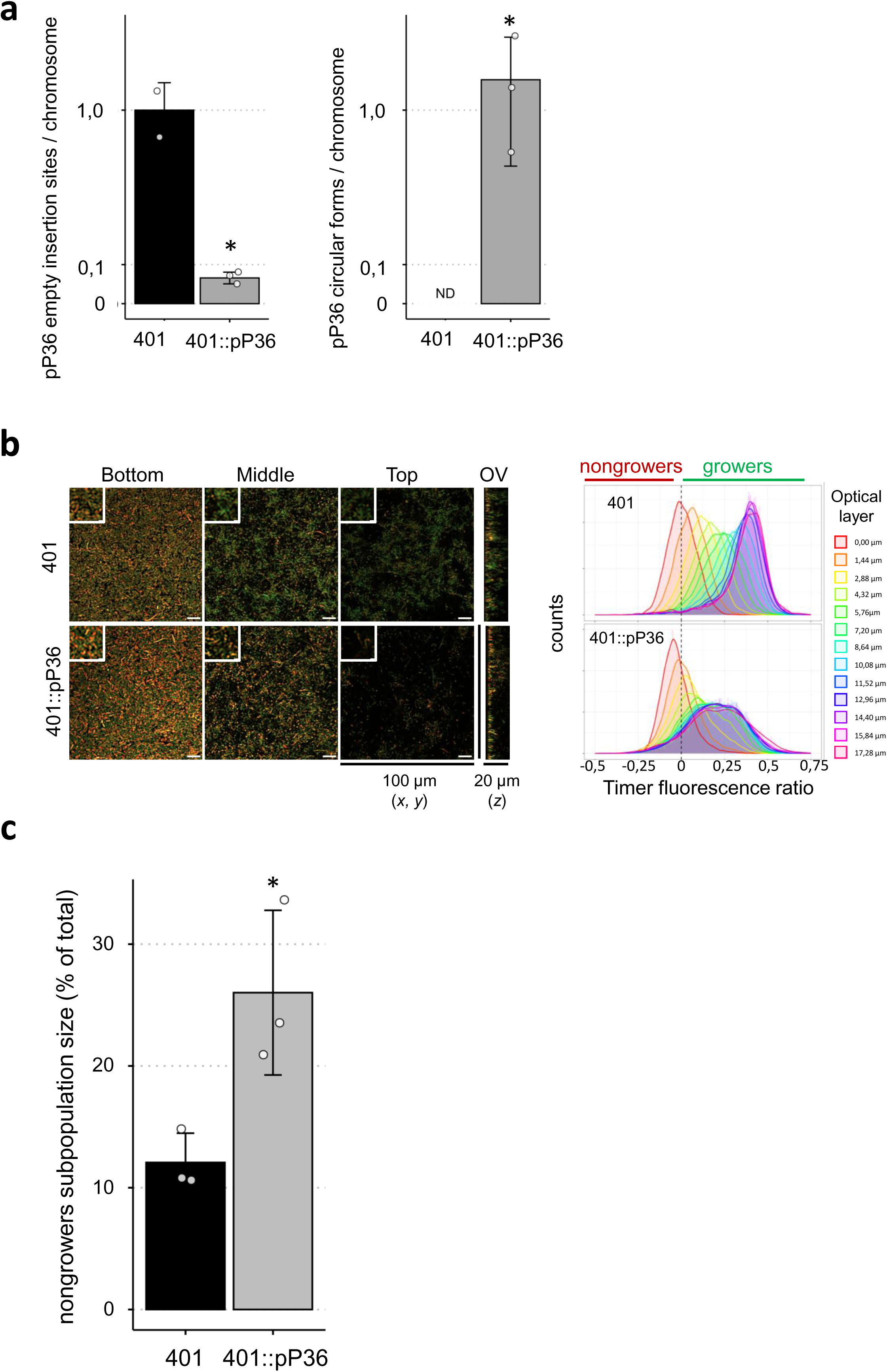
pP36 ICE mediates horizontal transfer of antibiotic persistence mechanisms. Linked to Figure 6. *L. pneumophila* strain 401 and *L. pneumophila* 401::pP36, producing Timer, were grown to stationary phase, diluted in AYE broth, allowed to form biofilms at 30 °C for 24 h. (**a**) qPCR analysis of pP36 empty sites and circular forms. Data represent the mean ± SD of three biological replicates (two-tailed Student’s t test; \*\*\**P* < 0.001, ** *P* < 0.01, \**P* < 0.05). (**b**) Confocal microscopy analysis. Micrographs (*x* = 100 μm, *y* = 100 μm) show overlays of the Timer fluorescence (500 nm and 600 nm); growing and nongrowing bacteria appear green or red/orange, respectively. OV: orthogonal view (*z* = 20 μm). Scale bars 10 μm. Distribution of the Timer fluorescence ratio calculated for each individual bacterium at different optical layer in the biofilm is displayed. (c) Quantification of the subpopulation size in nongrowers by confocal microscopy. Data represent the mean ± SD of a minimum of 3 biological replicates (two-tailed Student’s t test; \*\*\**P* < 0.001, ** *P* < 0.01, \**P* < 0.05).

## Supplementary information

Supplementary file: The *msevol* computational model of ICE dynamics

