## Supplementary information for "Heterogeneous dynamics of mobile genetic elements encode and spread antibiotic persistence in bacteria"

### The *msevol* computational model of Integrative and Conjugative Elements dynamics

#### Introduction

Mathematical models provide a valuable framework to explore the intricate dynamics of mobile genetic elements (MGEs) involved in the spread of antibiotic resistance. By incorporating interplayed processes such as horizontal gene transfer, selective pressures, and microbial population structure, they can help to improve our understanding of the mechanisms driving the persistence and evolution of resistance within microbial populations. In this context, we developed the ***msevol*** framework as a stochastic, multi-scale computational platform that integrates genetic, cellular, and ecological processes to simulate the emergence and maintenance of resistance under dynamic environmental conditions, providing a flexible tool to investigate how evolutionary and ecological factors jointly shape bacterial adaptation. We then used *msevol* to explicitly model the dynamics of integrative and conjugative elements (ICEs) which can excise from the chromosome and replicate intracellularly while imposing a metabolic cost to their host cells, allowing us to study how such mobile elements influence bacterial population persistence under selective pressure from antibiotics targeting actively dividing cells, even in the absence of resistance genes.

#### 1. Overview on the novel *msevol* computational framework

The *msevol* (Multi-Scaled EVOLution) framework was initially developed to simulate the stochastic dynamics of large bacterial populations in the context of antimicrobial resistance epidemics. Such biological systems are characterized by two key features: 1) a **nested organization of evolutionary units**, for example resistance genes carried by mobile genetic elements, which are themselves contained within bacteria organized in populations, which in turn may be structured across patients or hospital wards; and 2) **very large numbers of potentially identical elements**.

Nested systems (Figure SI 1A) are often represented using hierarchical "inclusion trees," in which each node of the tree represents a biological element and edges indicate inclusion relationships, linking genes to plasmids, plasmids to cells, and cells to populations (Figure SI 1B). While useful conceptually, this tree structure introduces redundancy: identical elements are duplicated each time they appear in a different context. In practice, this limits the size of populations that can be simulated and can bias the modeling of rare events, which are often critical for the emergence of antibiotic resistance. To overcome these limitations, *msevol* uses a compressed, non-redundant representation called a Minimal Directed Acyclic Graph (MDAG) (Figure SI 1C). In this representation, identical subtrees are merged and stored only once, which allows very large populations of similar elements to be handled efficiently. From a biological perspective, this means that stochastic events affecting many identical cells or genes can be computed once and applied across all equivalent elements, while still preserving the hierarchical structure of the system.

By combining the computational efficiency of population-level approaches with the flexibility of agent-based simulations, *msevol* can model large, structured bacterial populations and track the fate of MGE and/or resistance genes under different conditions. The computational complexity grows primarily with the number of unique biological entities and their interactions, rather than with the total population size, making it possible to explore rare but biologically important events in resistance evolution.

**Supplementary figure SI 1: Multiscale representations of the nested architecture of antimicrobial resistance.** Multiscale ecosystem models aim to represent biological objects (e.g. resistance genes, organisms, parasites and their hosts), as a system of nested structures (Panel A). A fundamental representation of this nesting hierarchy is a labelled tree T whose vertices represents instances of objects, vertex labels represent the category of the object and an edge represent the inclusion of one entity into another, such as a gene included in a chromosome in turn included in a bacterial cell (Panel B). However such a tree structure introduces redundancies affecting storage (e.g. the same 'red' resistance gene is stored four time) and execution time because evolution rules are applied to each individual instance of numerous, possibly identical objects (e.g. four individual Bernoulli trials to determine the occurrence of a mutation affecting the 'red' resistance genes), that induces inherent limitations in the simulation of very large populations. To circumvent such limitations, *msevol* is based on the manipulation of the multiplicity, minimal directed acyclic graph (MDAG) representation of the inclusion tree (Panel C). In MDAG representation, each unique instance (= same label and same content, or more formally, same induced subtree) is represented only once. For instance, the two 'purple' cells bacteria having both the same 'purple' chromosome the same 'grey' plasmid carrying the 'red' and 'blue' ARGs are represented as a single vertex, whose inner 'host -> cell' inclusion edge has a multiplicity of 2. On the contrary, the two 'orange' cells having the same label but different contents ('red' gene is carried either by the chromosome or by the plasmid) are still represented as two different vertices. In this example, the inclusion has size 55 (23 vertices and 22 edges) and its multiplicity MDAG equivalent has size 29 (13 vertices and 16 multiplicity edges). In the ecosystem simulation setting where multiplicities are typically very large (e.g. $10^9$ for bacterial cells), the minimal representation can lead to dramatic reductions of size requirements.

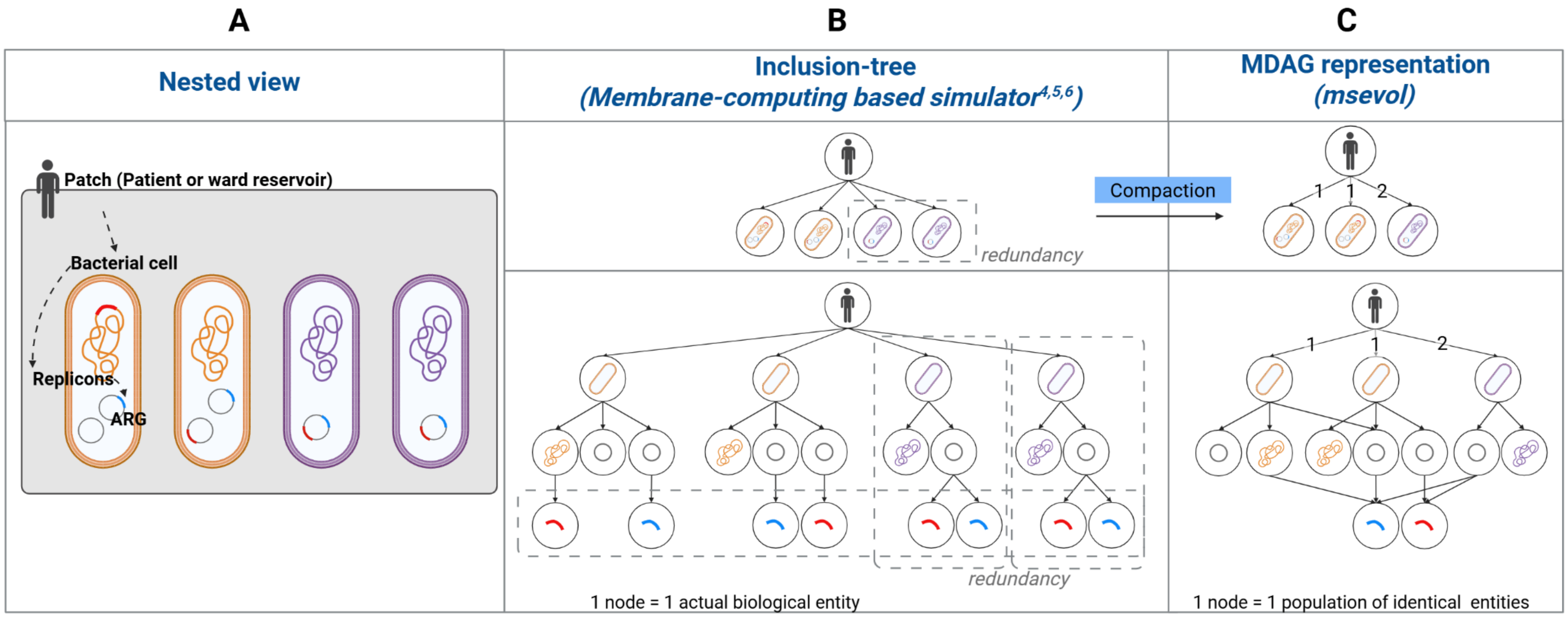

In practice, *msevol* operates through a dual architecture: a graph manipulation engine, defined independently of the simulated biological entities, coupled with a biological model tailored to the specific study. The biological model used in the present work is described below in Section 2, "ICE-Encoded Persistence Biological Model."

From an implementation perspective, the core *msevol* simulator, which encompasses the graph manipulation engine and the biological model, is written in C++. To enhance usability and facilitate data analysis, the software also includes *Rmsevol*, a companion R package. *Rmsevol* manages model input/output formatting, handles the execution of the C++ executable, and provides tools for data visualization and analysis. More details on the general principle of the MDAG manipulation used by *msevol* are available in the wiki of a preliminary version of the simulator (https://github.com/rasigadelab/msevol/wiki). The msevol simulator software, including both the C++ core engine and the applicative biological model dedicated to persistence carried by an ICE, can be freely downloaded from GitHub (https://github.com/rasigadelab/msevol2_funmge). It requires a Windows operating system (minimum Windows 10) for successful installation and compilation, along with Microsoft Visual Studio 2022 or later to build the C++ core executable. Similarly, the companion Rmsevol R package for this specific model, along with the code necessary to reproduce the simulations presented in this document and in the article, can also be downloaded from GitHub (https://github.com/rasigadelab/Rmse2FunMGE). It requires R Statistical Software (version 4.3.2 or later) to run simulations using Rmsevol.

The simulations were performed on a Windows 10 workstation equipped with an Intel® Xeon® CPU E5-2630 v4 (10 cores, 20 threads) running at 2.2 GHz, and 128 GB of RAM. The C++ core of the *msevol* simulator was compiled using Microsoft Visual Studio 2022, and the Rmsevol package was executed using R Statistical Software version 4.3.2. and additional R packages (*"data.table"* version 1.15.0; *"magrittr"* version 2.03); "*visNetwork*" version 2.1.2; *"ggplot2"* version 3.4.4; *"cowplot*" version 1.1.3; *"ggpubr*" version 0.6.0; and "*segmented*" version 2.1.4). All simulations were run under default system settings, without parallelization, to ensure reproducibility.

#### 2. ICE-encoded persistence biological model

The biological entities included in the present applicative biological model are summarized in Supplementary Figure SI 2 and supplementary table SI 1**.** Briefly, this model focuses on the dynamics of an integrative and conjugative element (ICEs), which can excise from the bacterial chromosome, replicate intracellularly, and impose a metabolic cost on their host cell. Bacterial cells containing chromosomic or free ICEs are structured into metapopulations (also called patches), where they compete for resources under shared antimicrobial stress. Cells can diversify from their initial genotype through ICE excision from the chromosome, after which ICEs can undergo stochastic replication or loss. Upon excision, ICEs may become metabolically costly for the host cell, with the total cost increasing with the number of free ICE copies, creating variability in cell fitness across the population. Antibiotic induced mortality is assumed to be growth-dependent, meaning that cells These eco-evolutive dynamics are simulated through the sequential application of 5 discrete events (Supplementary Figure SI 2, Panel C).

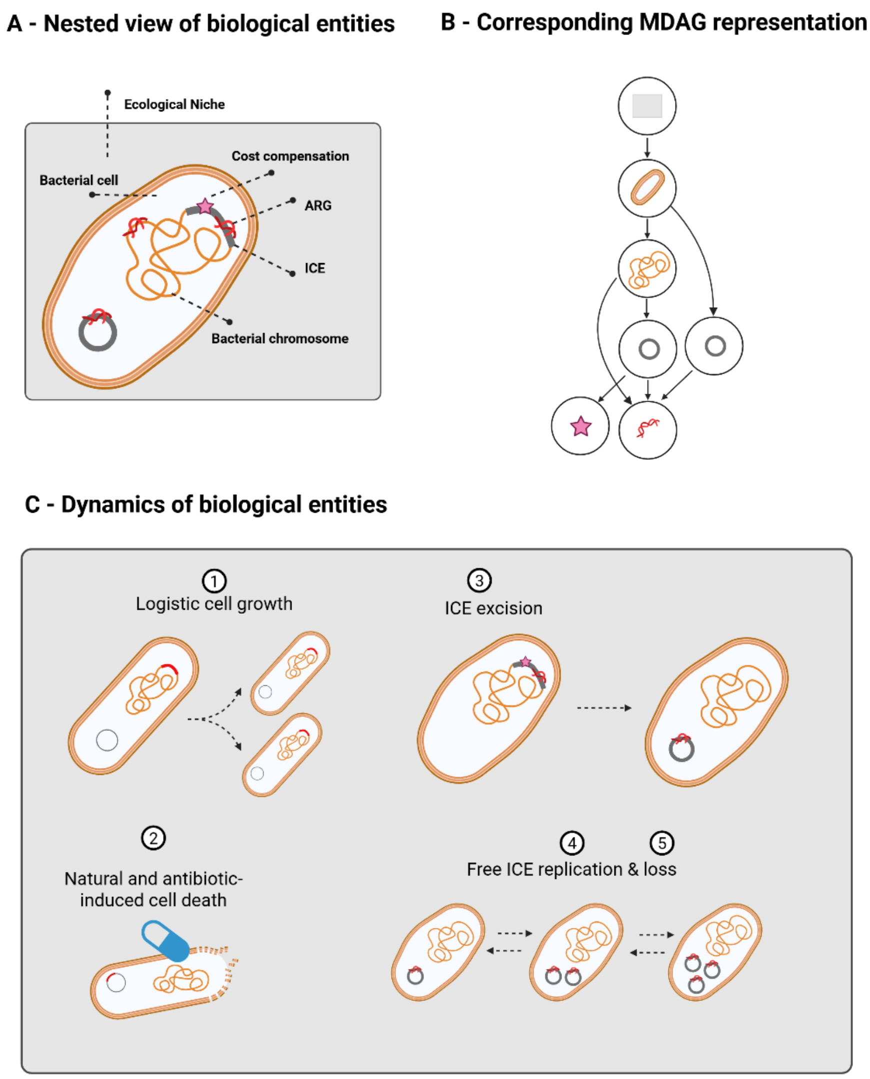


**Supplementary Figure SI 2: Applicative model.**

Six types of agents are considered in the current model. Patches represent a set of interconnected bacterial metapopulations in which bacterial cells compete for shared resources and are exposed to the same antimicrobial selective pressure. Bacterial cells within these patches carry various replicons, which can be either chromosomes or ICEs. Replicons contain antibiotic resistance genes (ARGs) that reduce the sensitivity of their bacterial host to bactericidal antimicrobials. ICEs can be inserted into chromosomes or exist as free, circular elements in the cytoplasm. They may also contain an artificial agent, costCompensation, introduced to modulate the fitness cost of the ICE depending on its state. Notably, this cost compensation is lost by the ICE upon excision, whereas all other ICE contents (e.g. ARGs; see Panel C, rule 3) are retained.

**Supplementary table SI 1: Biological entities and their properties included in the biological msevol model**. Intrinsic properties correspond to the fixed, user-defined characteristics of a biological agent; they are defined independently of the agent's contents. Current properties correspond to the effective characteristics of the agent, i.e. they result from the modulation of the agent's intrinsic properties by its direct descendants. For instance, a bacterium carrying a costly ICE actually grows at a reduced effective growth rate, whereas a cell without the ICE grows at an effective rate equal to its intrinsic growth rate. These current properties are automatically computed by *msevol* whenever the contents are modified.

| Biological agent | Intrinsic properties | Current properties |
|---|---|---|
| **Ecological niche** | Carrying capacity K<br>Selective pressure $k_{kill}$ | |
| **Bacterial cell** | Intrinsic growth $\beta_0$<br>Intrinsic natural death rate $\delta_0$<br>Intrinsic resistance $R_0$<br>Growth-killing coupling factor $\alpha$ | Current "growth"<br>$\beta^{eff} = \beta_0 + \sum_{i=cell's\ children} \beta_i^{eff}$<br>Current resistance<br>$R^{eff} = R_0 + \sum_{i=cell's\ children} R_i^{eff}$ |
| **Chromosome** | Intrinsic cost/benefit $\beta_{chrom}^0$<br>Intrinsic resistance $R_{chrom}^0$ | Current "growth"<br>$\beta_{chrom}^{eff} = \beta_{chrom}^0 + \sum_{i=chrom\ children} \beta_i^{eff}$<br>Current resistance<br>$R_{chrom}^{eff} = R_{chrom}^0 + \sum_{i=chrom.'s\ children} R^{eff}\ _i$ |
| **Integrative and conjugative element (ICE)** | Intrinsic cost/benefit $\beta_{ICE}^0$<br>Intrinsic resistance $R_{ICE}^0$<br>Excision rate $r_{exc}$<br>Replication rate $r_{rep}$<br>Loss rate $r_{loss}$ | Current "growth"<br>$\beta_{ICE}^{eff} = \beta_{ICE}^0 + \sum_{i=ICE\ children} \beta_i^{eff}$<br>Current resistance<br>$R_{ICE}^{eff} = R_{ICE}^0 + \sum_{i=ICE\ children} R_i^{eff}$ |
| **CostCompensation** | Intrinsic cost/benefit $\beta_{comp.}^0$ | Current "growth"<br>$\beta_{comp.}^{eff} = \beta_{comp.}^0$<br>*(no possible direct content)* |
| **ARG (not used)** | Intrinsic resistance $R_{ARG}^0$ | Current resistance $R_{ARG}^{eff} = R_{ARG}^0$ |

##### 2.1. Cell dynamics events

**[Event 1] Cell growth**

The growth of a cell in a given patch follows a logistic framework, meaning that the rate of division decreases as the patch becomes crowded, reflecting competition for limited resources. The division rate ranges from a maximum rate (referred to as $\beta^{eff}$), when the patch is empty, down to zero as the patch occupancy approaches its carrying capacity $K$. The value of $\beta^{eff}$ is itself determined from the cell's basal growth rate $\beta_{basal}^0$, which represents its fixed, intrinsic potential to divide, and is reduced (respectively increased) by the metabolic costs (respectively benefits) associated with any mobile genetic elements $i$ it carries, such as ICEs, so that cells with costly elements divide more slowly even at low population density:

$$\beta_{birth,j,k} = \beta_{eff,j} \underbrace{\left(1 - \frac{N_k}{K}\right)}_{logistic\ penalization} N_j$$

$$= \underbrace{\left(\beta_{basal,j} + \sum_{i\,\in\{direct\ content\ of\ cell\ j\}} \beta_{eff,i}\right)}_{Additive\ modulation\ of\ cell's\ intrinsic\ fitness\ by\ the\ "fitness"\ of\ its\ contents} \left(1 - \frac{N_k}{K}\right) N_j$$

At each time step, the number of identical cells $j$ that actually divide in patch $k$ is drawn randomly from a binomial distribution, where the probability of division for each cell is derived from its growth rate using the approximation $P_{birth,j,k} \approx 1 - e^{-\beta_{birth,j,k}\,\Delta t}$, linking the continuous growth rate to a discrete event. Daughter cells are assumed identical to their ancestor cell, including notably the same number of ICEs (no random partitioning of mobile genetic elements during division).

**Supplementary Figure SI 3: Simple logistic growth**. [Left] The simulation was initiated with a single bacterial population growing at a maximal, intrinsic rate of 0.5 per hour in a closed environment without antibiotic pressure. [Right] The corresponding time–cell dynamics simulated by msevol are shown.

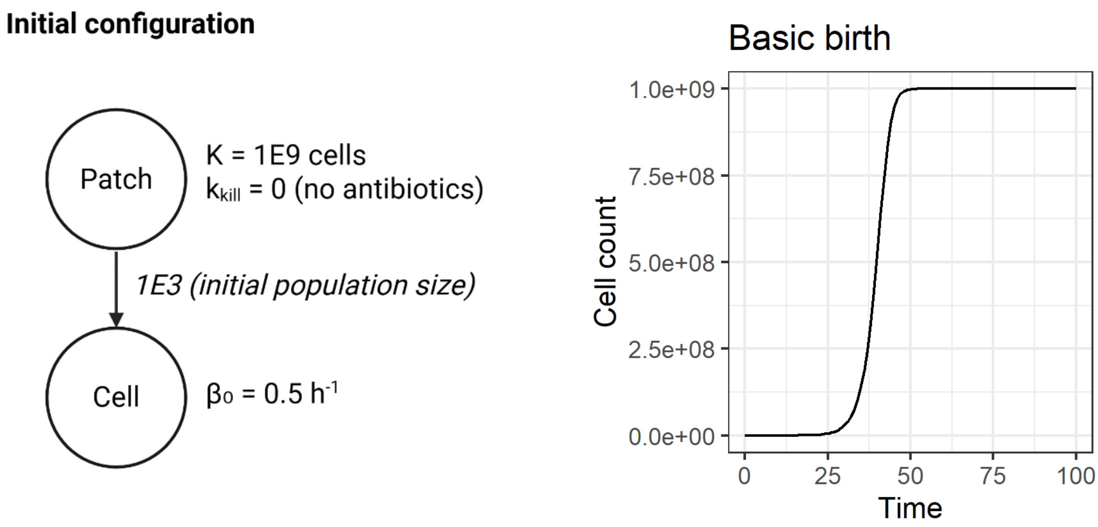

**Figure SI 4: Same logistic cell growth with penalized fitness**

The figure illustrates the transitivity of the fitness growth penalty imposed by resistance genes and/or mobile genetic elements (in this case, integrative and conjugative elements, ICEs) on their host cells. The scenario simulates independently the growth of four cells with the same intrinsic growth rate ($\beta_0$ = 0.5 per hour) but differing in their ICE content. (1 -Black curve) The first cell is ICE-free (no growth penalty); (2- blue curve) the second cell carries a costly chromosomal ICE, imposing a fitness cost of −0.2 per time step; (3 - green curve) the third cell carries the same chromosomal ICE, but this ICE also contains a “CostCompensation” element that exactly offsets the cost (i.e. confers a fitness modulation of +0.2 per time step), thereby neutralizing the apparent cost of the ICE; and (4 – red curve) the final cell carries two copies of the same costly ICE (one chromosomal and one extrachromosomal), resulting in a doubled total cost. Cells are assumed to grow independently in their own ecological niches (i.e. independent simulations with no intercellular competition).

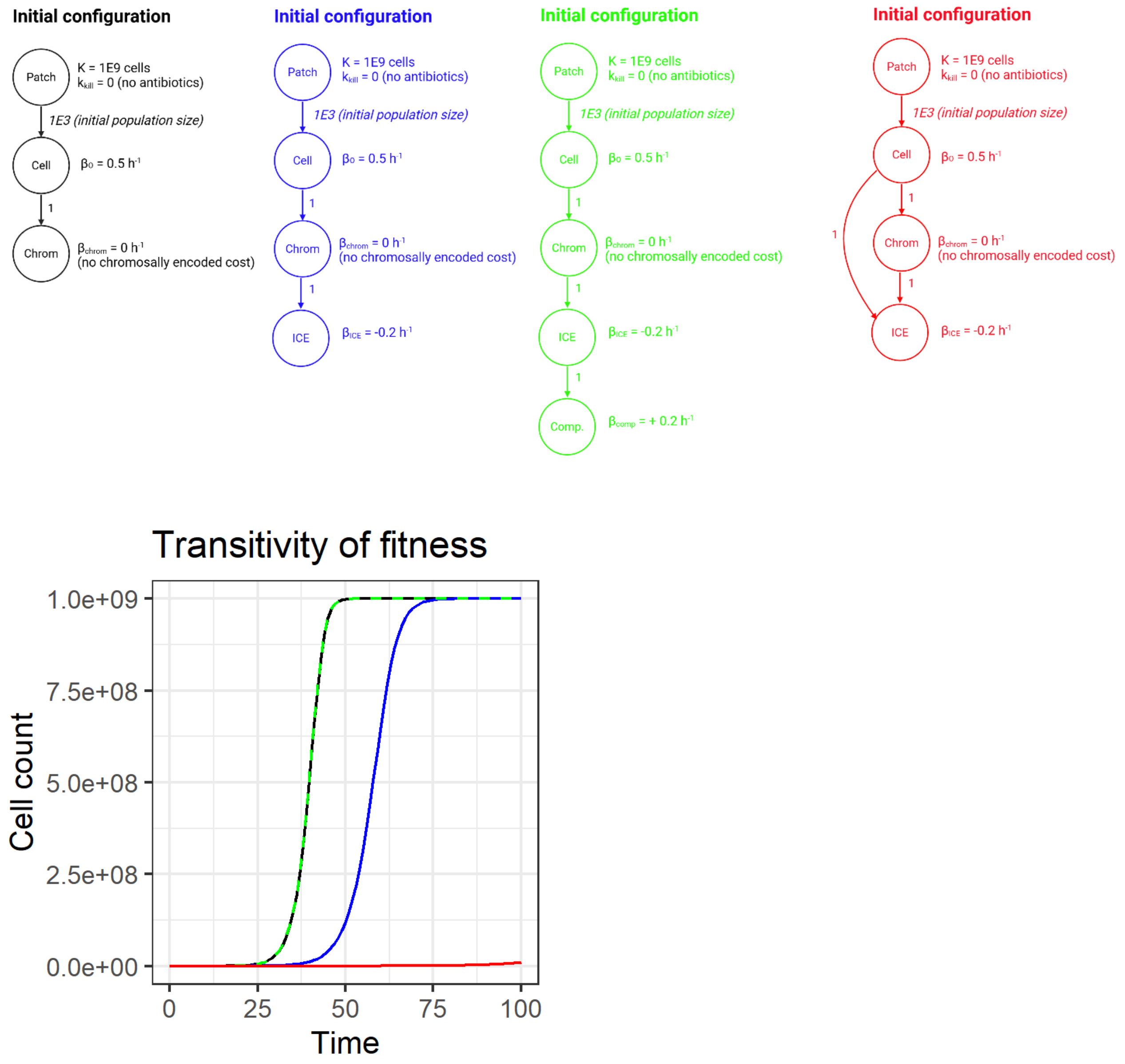

**[Event 2] Cell death (natural and antibiotic-killing induced)**

The mortality of a cell in a given patch (i.e. ecological niche) is described by an effective death rate $\delta_j$, which integrates both intrinsic (natural) and antibiotic-induced sources of mortality. The natural component of mortality ($\delta_{0,j}$) represents the baseline death rate in the absence of treatment, arising from factors such as aging, metabolic stress, or stochastic physiological failure. In the presence of an antibiotic, an additional killing rate ($k_{kill}$) contributes to mortality, reflecting the action on fully sensitive, actively dividing cells. This antibiotic-mediated mortality is further modulated by two key cellular properties:

- **Resistance mechanisms**, quantified by a resistance level R, alter the effective killing rate, with higher resistance values reducing antibiotic efficacy according to an exponential factor $2^{-R}$. Such a formulation is conceptually analogous to a minimum inhibitory concentration (MIC) scale, since each unit increase in R corresponds to a twofold decrease in susceptibility, that is, to a doubling of the MIC value.

- **Growth-related modulations**, account for the fact that some antibiotics may preferentially targets actively dividing cells. In the absence of more precise information, the dependency of antibiotic killing on growth activity is assumed to be linear and expressed as
$$(1-\alpha)+\frac{\beta_{eff,j}}{\beta_{0,j}}*\alpha$$
where $\beta_{eff,j}$ represents the effective growth rate of the cell relative to its intrinsic (reference) growth rate $\beta_{0,j}$. The parameter $\alpha$ (ranging from 0 to 1) determines the strength of coupling between antibiotic killing and growth: when α=1, non-dividing cells are fully protected from antibiotics, whereas when α=0, mortality becomes completely independent of growth activity.

Combining these contributions, the effective mortality rate of cell j can be expressed as a single equation integrating natural death, antibiotic-induced killing, resistance, and growth-dependent modulation:

$$\delta_{j,k} = \underbrace{\delta_{0,j}}_{Natural\ death\ rate} + \underbrace{k_{kill,k}}_{Antibiotic-induced\ killing\ rate} * \underbrace{\left[2^{R_j}\right]}_{Modulation\ by\ resistance} * \underbrace{\left[(1-\alpha)+\frac{\beta_{eff,j}}{\beta_0}*\alpha\right]}_{Modulation\ \ by\ effective\ growth}$$

Like for growth event, the mortality event is applied to the population of $N_{j,k}$ identical cells of type j contained in patch k. The number of cells that actually die is then drawn randomly from a binomial distribution, where the probability of death for each cell is derived from its effective mortality rate using the approximation $P_{birth,j,k} \approx 1 - e^{-\delta_j\,\Delta t}$, linking the continuous death rate to a discrete stochastic event.

**Figure SI 5: Growth-dependent antibiotic-induced killing**

[Top] The panel illustrates the shape of the growth modulation factor induced by a penalized growth rate (i.e. the shape of $\underbrace{\left[(1-\alpha)+\frac{\beta_{eff,j}}{\beta_0}*\alpha\right]}_{Modulation\ \ by\ effective\ growth}$ ). Cells growing at an effective rate equal to their intrinsic rate (i.e. with no cost penalization from their contents) or at a higher rate (i.e. when their contents are beneficial) experienced maximal killing whatever the value of the growth-killing parameter α. On the contrary, non-growing cells experience minimal killing, i.e. the fraction of the killing that is independent to growth (= 1- α).

[Bottom] The panel shows the dynamics of three cells with the same intrinsic growth rate ($\beta_0$ = 0.5 per time step) but differing in their ICE content and consequently in their growth dependence for antibiotic killing. (Black curve) ICE-free cell (no growth penalty), with α = 1. Killing is maximal since there is no growth reduction from the basal rate; (Red curve) Cell carrying a costly ICE (growth penalty of −0.3 per time step), with α = 1. Killing is reduced, leading to prolonged persistence of the red cell population. (Green curve) Cell carrying the same costly ICE (growth penalty of −0.3 per time step), but with a lower α = 0.5, meaning a weaker coupling between growth and killing. However, the killing reduction is insufficient to offset the growth cost, resulting in a lower net growth rate and thus a shorter persistence of the green cell population. Cells are assumed to evolve independently in their own ecological niches (i.e. independent simulations with no intercellular competition), under constant selective pressure ($k_{kill}$ = 1 per unit time, corresponding to approximately 2 times the stress that compensates the growth of fully sensitive cells).

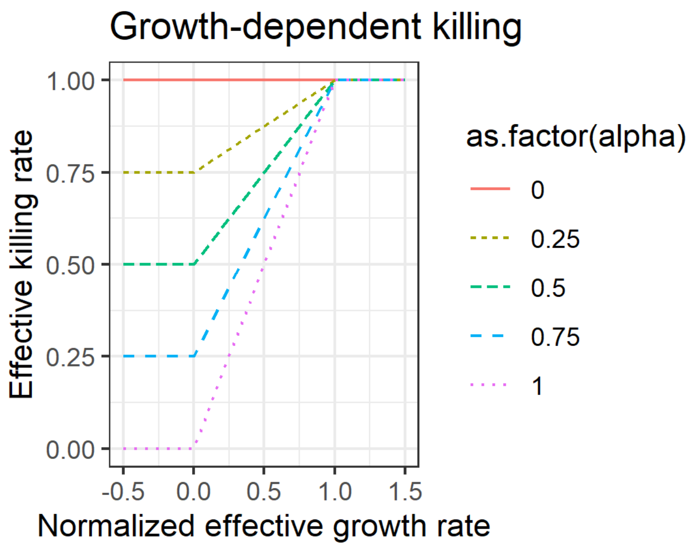


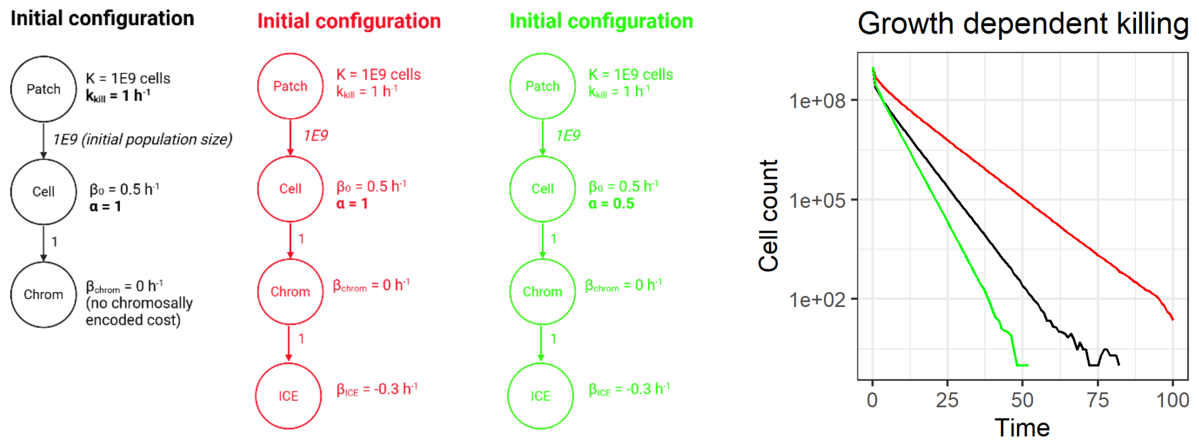

#### 2.2. Within-cell ICE dynamics events

**[Event 3] Excision of chromosomal ICE**
ICEs can excise from the host chromosome to form free, autonomously replicating DNA molecules capable of subsequent replication or horizontal transfer. Excision is catalyzed by ICE-encoded recombinases and often regulated by host factors, environmental signals, stress responses, or cell-cycle cues. Some ICEs impose a metabolic cost on their host, which may differ depending on whether the ICE is integrated in the chromosome or exists as a free copy. For example, an ICE may be largely silent and impose little or no cost when integrated, but its autonomous replication or expression of mobilization functions after excision can increase the metabolic burden on the cell.

In the *msevol* model, excision is implemented as a stochastic event with a constant, intrinsic excision rate $r_{excision,i}$., independent of host context. For a cell carrying $N_{i\ in\ j}$ copies of chromosomally integrated ICE$i$ (typically equal to 1), the actual number of copies excised during a simulation time step $\Delta t$ is drawn from a binomial distribution with parameters $N_{i\ in\ j}$ and $P_{excision,i} \approx 1 - e^{-r_{excision,i}\,\Delta t}$.

To account for the differential metabolic cost of ICEs depending on their state, we introduced a dedicated entity, **CostCompensation**, which can fully or partially offset the metabolic burden of the ICE when integrated in the chromosome. Upon excision, the ICE is assumed to lose all its compensations, so the host cell immediately experiences the full metabolic cost of carrying the free ICE. This mimics situations where an ICE is largely unexpressed and non-costly when integrated, but becomes metabolically burdensome once excised.

**[Events 3] Free ICE replication**
The intracellular replication of free integrative and conjugative elements (ICEs) within a host cell is described by an effective replication rate $r_{replication,i}$ (where the index $i$ refers to the ICE), reflecting the element's intrinsic ability to duplicate autonomously. Biologically, this captures the ability of an ICE to replicate, once excise from the chromosome, as an independent DNA molecule prior to reintegration or conjugative transfer. As for previous events, the actual number of ICE copies that replicate in a host cell is drawn randomly from a binomial distribution with parameters $N_{i\ in\ j}$ (the number of ICE copies in the cell) and $P_{rep,i} \approx 1 - e^{-r_{replication,i}\,\Delta t}$(the probability of replication per copy).

**[Events 4] Free ICE loss**
Loss of free ICEs can occur through several mechanisms, including segregational loss during host cell division, or stress- or damage-induced DNA degradation. In our *msevol* model, these processes are captured by a stochastic loss term defined by a constant effective loss rate $r_{loss,i}$. The actual number of ICE copies lost in a host cell during a simulation time step is drawn from a binomial distribution with parameters $N_{i\ in\ j}$ (the number of ICE copies in the cell) and $P_{rep,i} \approx 1 - e^{-r_{replication,i}\,\Delta t}$(the probability of loss per copy).

**<u>Note</u>** that the dynamic parameters governing the three ICE events described above are all assumed to be intrinsic properties of the ICE and independent of host context, in order to maintain a mechanistic yet simple modeling framework in the absence of further biological characterization. Nevertheless, the overall persistence and propagation of an ICE in the population also depends on vertical transmission through host cell growth and division, which is modulated by any fitness costs associated with carrying the element, including the differential metabolic burden between the integrated and excised states.

**Figure SI 6: Excision of chromosomal ICEs, with loss of their putative "CompensationCost"**
[Top left] Initial model graph configuration.
[Top right] Model graph after one-time step. The ecosystem now contains two distinct cell populations with the same intrinsic properties but different genome architectures: one population contains a chromosomal copy of the ICE, and the newly created population contains a free copy of the ICE. Note that the loss of the "costCompensation" agent during excision led to the creation of a new "ICE" node in the graph. Both cells (respectively both ICEs) share identical intrinsic properties but differ in effective growth rate due to the loss of the "costCompensation" agent, which transitively modifies the effective "growth" of the ICE and, consequently, of the host cell.
[Bottom right] Time dynamics of the number of cells according to their effective birth rate. These dynamics reflect the excision process.
[Bottom left] Time dynamics of the total number of ICEs (black) and the total number of free, circular ICEs (red curve).

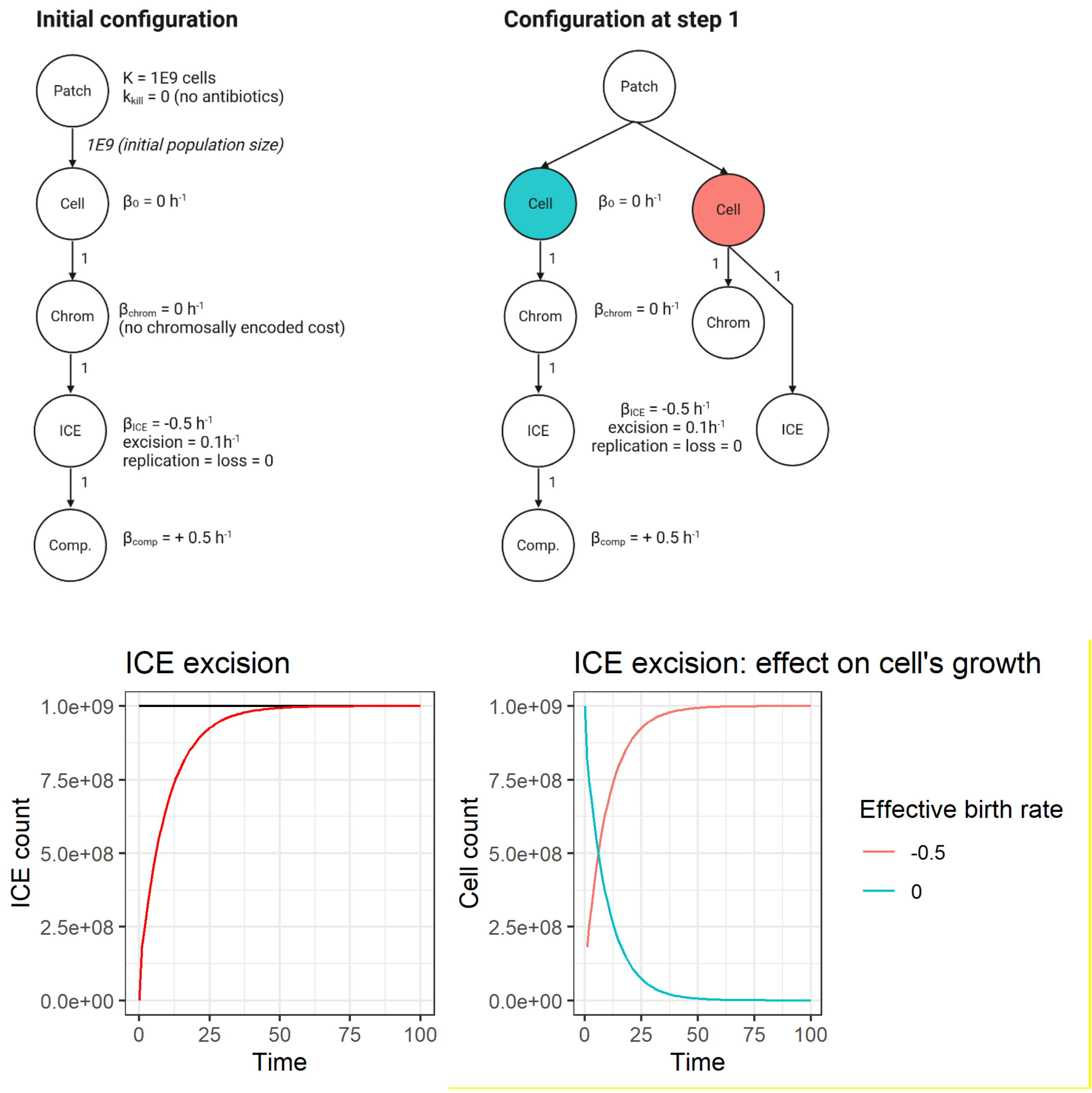

**Figure SI 7: free ICE intracellular replication**

[Top left] Initial model graph configuration. [Top right] Model graph after two time steps. The ecosystem now contains distinct cell populations with the same intrinsic properties but differing in the copy number of free ICEs.[Middle] Time dynamics of the total number of ICEs in the ecosystem. This number increases over time as a result of random intracellular replication of free ICEs.[Bottom] Cell diversification, recorded by their effective growth rate (which corresponds to their number of ICEs), resulting from intracellular ICE replication. Each panel represents the full distribution of cells at a single time point (indicated by the panel title).

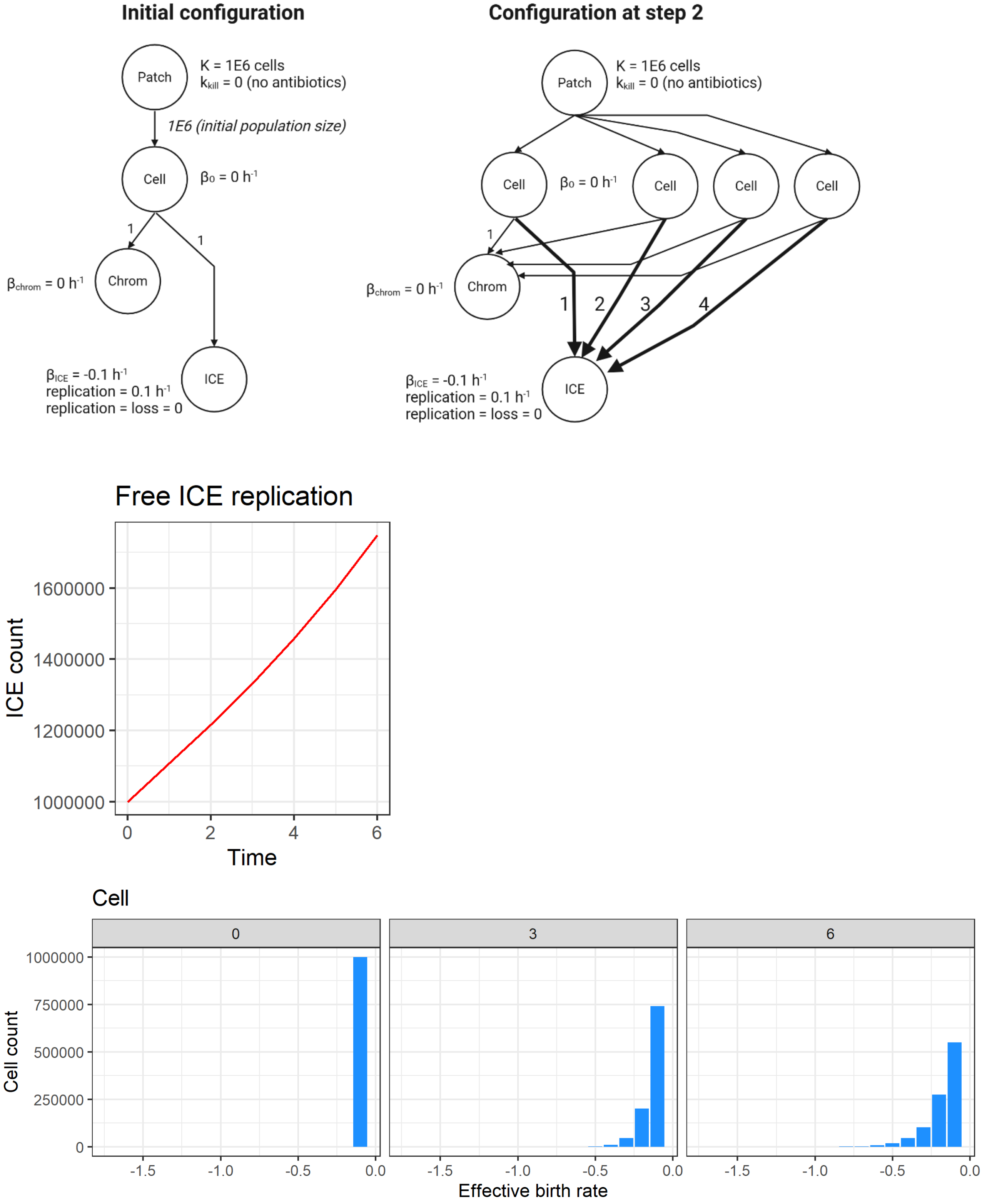

#### 3. Exploratory simulations

We used *msevol* to explore different sources of bacterial growth heterogeneity and their impact on time-kill dynamics, as presented in the main text.

**Supplementary table SI 2: Baseline parameters used for the simulations**

| Parameter | Value | Reference |
|---|---|---|
| Initial population size | $10^6$ cells | This simulation |
| Carrying capacity | $10^{12}$ cells | This simulation. Set high enough to prevent from logistic growth saturation effect. |
| Intrinsic growth rate | 0.2 per hour, corresponding to a doubling time of ~3.5h | Value defined according to the Timer fluorescence ratio correlation with Legionella pneumophila division rate [1] |
| Cost of the free ICE | -0.15 per hour, leading to a growth reduction of 75% | This study |
| Free ICE excision rate | 0.01 per hour | This study. Value compatible with switch rate estimated by other modeling approaches [2] |
| Free ICE replication rate | 0.01 per hour | This study |
| Free ICE loss rate | 0.001 per hour | This study |
| Antibiotic killing rate | 0.6 per hour | This study.<br>Lead to 99% cell elimination in ~11h and 99.99% in ~23h for fully sensitive cells. |
| Growth-killing coupling factor | 0.9 | This study. |

##### 3.1. Emergence of biphasic time-kill curve from fixed growth heterogeneity

We first explored how biphasicity may arise from fixed, chromosomally encoded growth heterogeneity. The population was divided into subpopulations with predetermined growth rates, either as two well-separated groups or as multiple subpopulations approximating natural variation. Biphasicity emerged in the two-subpopulation scenario under strong growth–killing coupling and a sufficiently large fraction of slow-growing cells (see main text, Figure 1B), whereas populations with normally distributed growth variations rarely displayed biphasic behavior (Figures SI 7 and SI 8 below).

**Figure SI 8: Initial setup of the simulations mimicking natural variations of growth**

[Top] This model represents a time–kill experiment involving J bacterial subpopulations evolving in competition in a closed system under constant antibiotic selective pressure. All subpopulations are assumed to be derived from the same basal cell line but exhibit a chromosomally encoded, fixed variation in growth rate. The populations undergo competitive growth and antibiotic-mediated killing, leading to changes in cell numbers over time, while cells remain unable to diversify from their initial genotype (notably, daughter cells inherit their mother's growth rate).

[Bottom] The effective growth rates of each subpopulation and the initial subpopulation sizes are automatically computed from a discretization of the normal distribution with a mean equal to the intrinsic growth rate and a user-specified standard deviation, mimicking natural growth variation. This includes subpopulations with reduced growth as well as subpopulations with enhanced growth relative to the intrinsic value. Subpopulations are assigned growth rates via discretization of a normal distribution with mean μ (user-defined intrinsic growth rate) and standard deviation σ (user-defined, representing natural variability in growth among cells within a clonal population). The J subpopulations are defined as follows: J–2 intermediate categories evenly span the interval [μ – x·σ, μ + x·σ] (with x user-defined, typically set to 3 to cover ~99.7% of the population), while the remaining two categories cover the lower and upper extremes. For each interval, the representative growth rate is calculated as the truncated mean. Initial subpopulation sizes are then randomly drawn from a multinomial distribution with probabilities corresponding to these intervals, ensuring that the population collectively reflects the prescribed growth variation. The figure illustrates the procedure for an intrinsic growth rate of 0.2 $h^{-1}$, with σ = 0.05 $h^{-1}$ (~25%) and J = 7 subpopulations. The blue curve represents the continuous normal distribution, the red dots indicate the representative growth rates of each subpopulation (equal to intrinsic growth rate + $cost_j$), and the histogram shows the normalized size of each subpopulation.

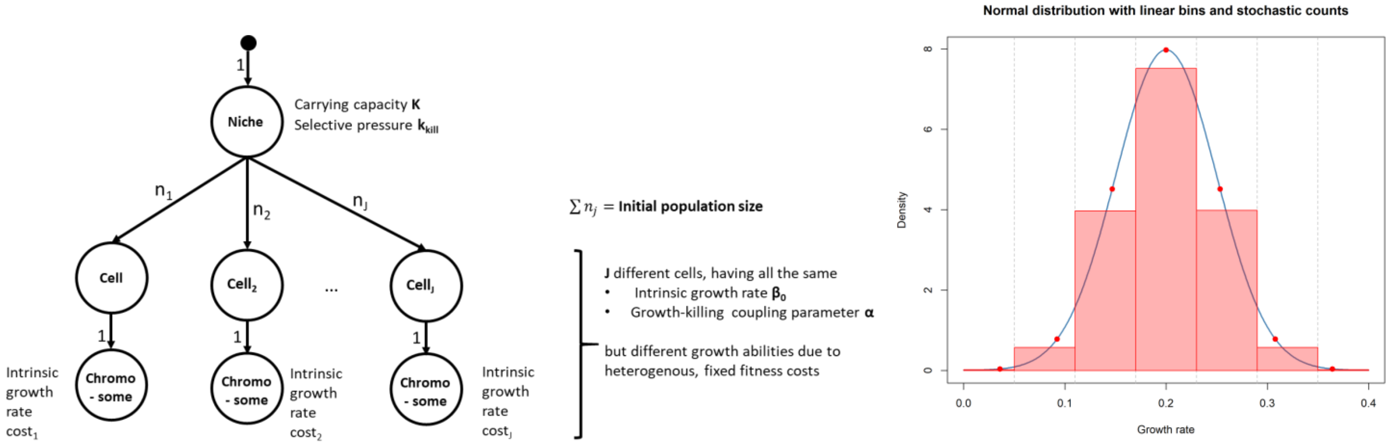

**Figure SI 9: Simulated time-killing assay of a cell population with natural variation of growth relative to their intrinsic grow rate, according to various initial settings.** The curves represent the dynamics of the total cell population, averaged over N = 20 simulations. Colors indicate the number of subpopulations (7 or 11), which has no noticeable effect on the overall dynamics. Each panel corresponds to a combination of α (rows) and assumed growth variability (columns). Biphasic dynamics may emerge at high α and high natural variation, suggesting that natural variability alone may be less effective at generating persistence than maintaining two distinct populations.

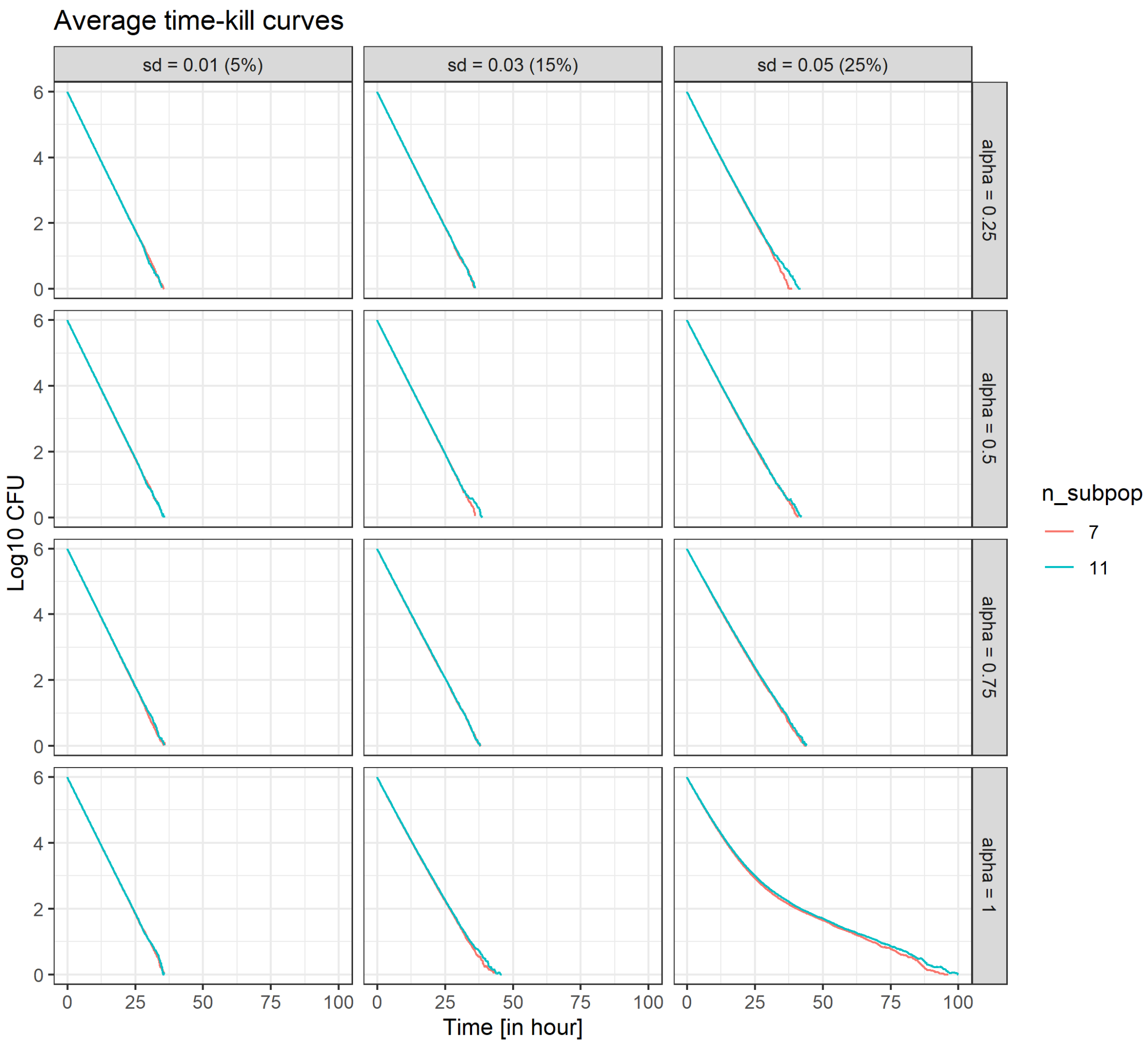

##### 3.2. Emergence of biphasic time-kill curve from growth heterogeneity induced by ICE dynamics

We then explored how biphasic time-kill curves can emerge from growth heterogeneity induced by ICE dynamics. In this framework, a fraction of cells randomly undergoes abrupt growth reduction upon ICE excision, producing subpopulations with distinct, stable growth rates. Allowing excised ICE to replicate or be lost dynamically generates a continuum of growth rates, creating more graded and heterogeneous growth behavior across the population. In both cases, biphasic killing curves arise under strong growth–killing coupling, with persistence prolonged by the presence of slow- or non-growing subpopulations, illustrating how ICE dynamics can drive population-level survival under antibiotic pressure.

**Figure SI 10: Simulated time–kill assay of an initially homogeneous bacterial population carrying an integrated ICE, according to various cost, excision and duplication rates.** Curves represents the dynamics of the total cell population, averaged over N = 20 simulations.

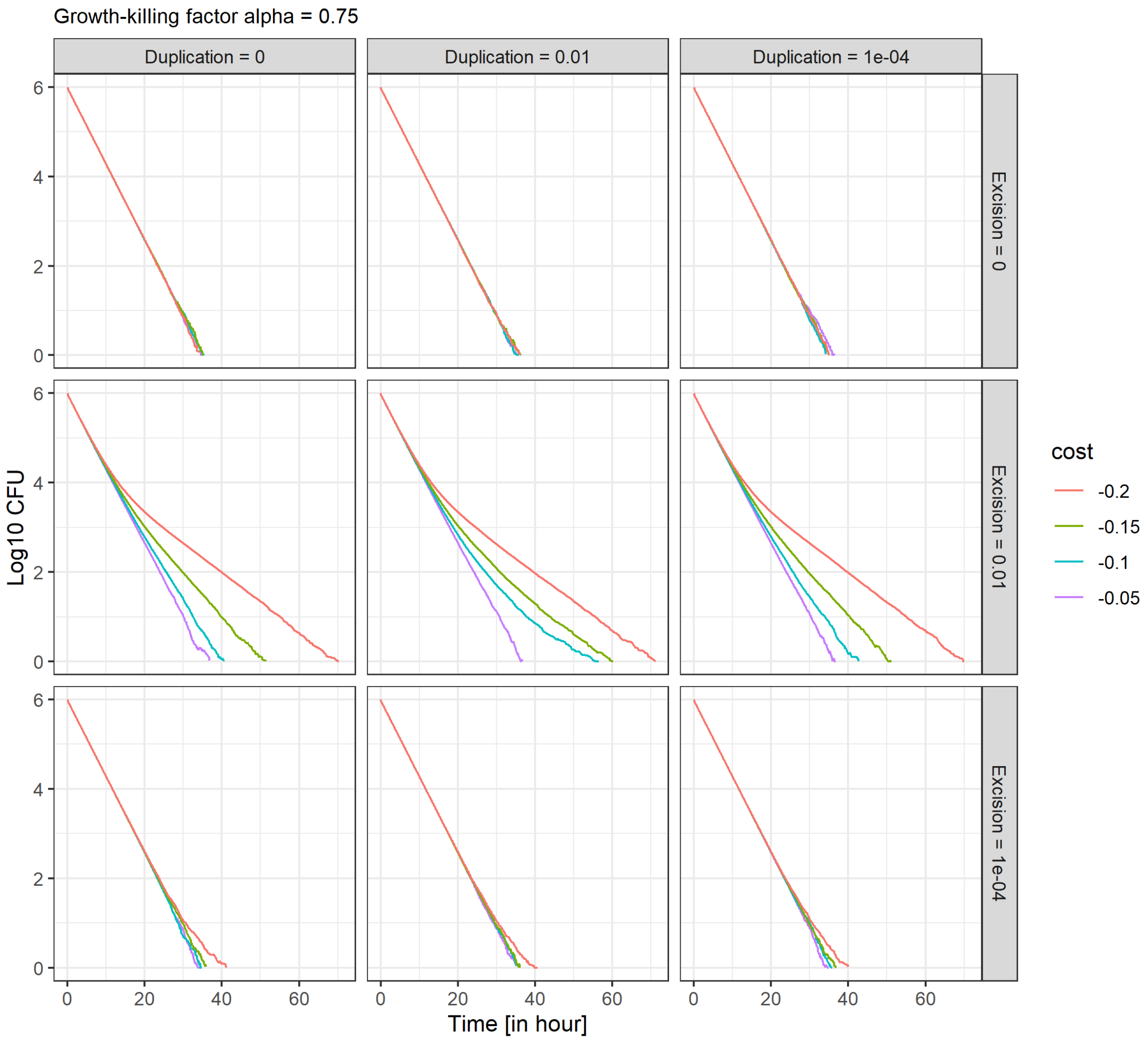

**Figure SI 11: Same as SI8 for a higher growth-killing coupling factor**

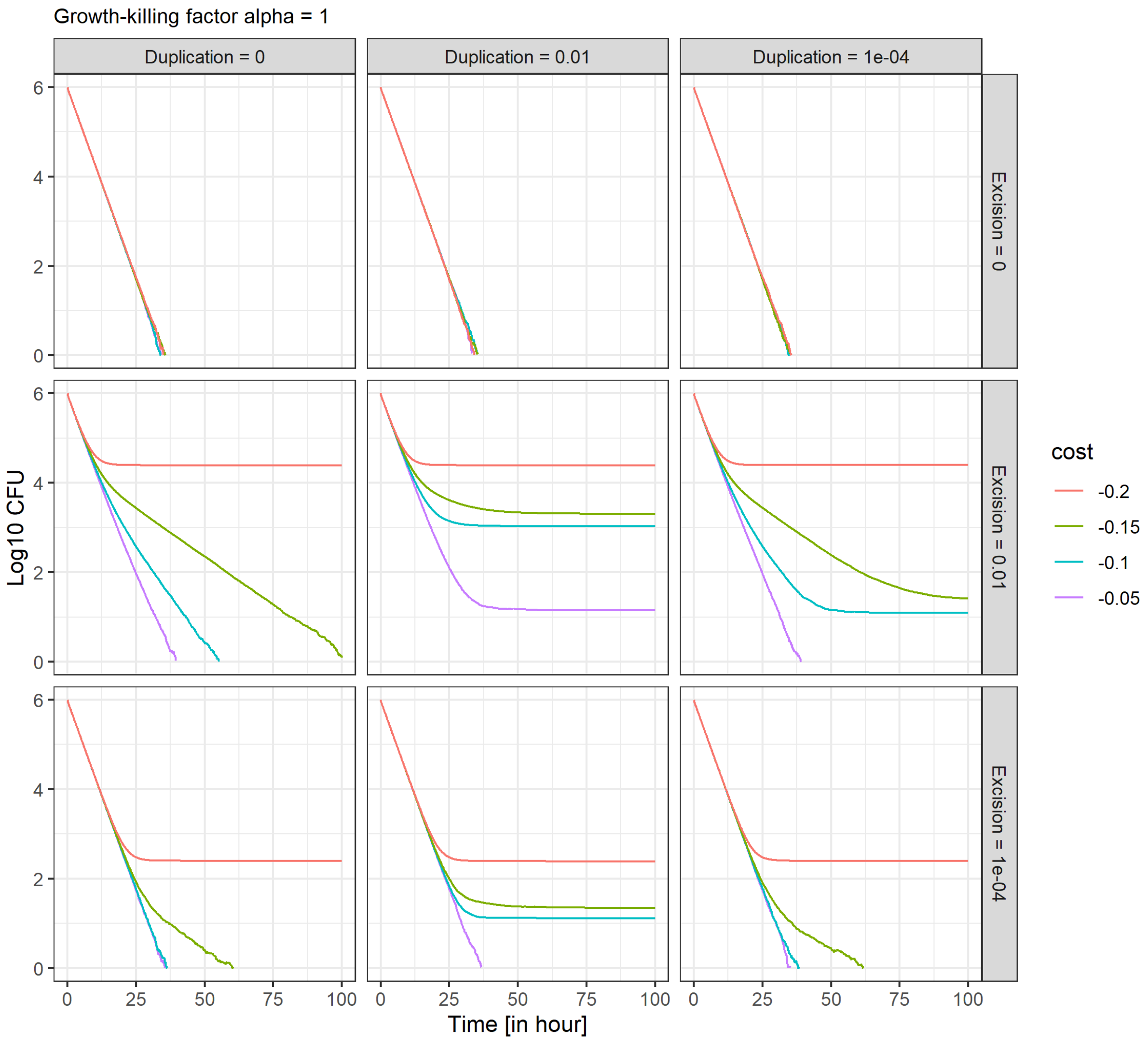


#### Supplementary information references


1 Personnic, N. *et al.* Quorum sensing modulates the formation of virulent *Legionella* persisters within infected cells. *Nature communications* **10**, 5216 (2019). https://doi.org:10.1038/s41467-019-13021-8

2 Carvalho, G., Guilhen, C., Balestrino, D., Forestier, C. & Mathias, J. D. Relating switching rates between normal and persister cells to substrate and antibiotic concentrations: a mathematical modelling approach supported by experiments. *Microb Biotechnol* **10**, 1616-1627 (2017). https://doi.org:10.1111/1751-7915.12739
